# Transient regulation of focal adhesion via Tensin3 is required for nascent oligodendrocyte differentiation

**DOI:** 10.1101/2022.02.25.481980

**Authors:** Emeric Merour, Hatem Hmidan, Corentine Marie, Pierre-Henri Helou, Haiyang Lu, Antoine Potel, Jean-Baptiste Hure, Adrien Clavairoly, Yi Ping Shih, Salman Goudarzi, Sebastien Dussaud, Philippe Ravassard, Sassan Hafizi, Su Hao Lo, Bassem A. Hassan, Carlos Parras

**Affiliations:** Paris Brain Institute, Sorbonne Université, Inserm U1127, CNRS UMR 7225, Hôpital Pitié-Salpêtrière, 75013 Paris, France; Department of Biochemistry and Molecular Medicine, University of California-Davis, Sacramento, CA 95817, USA; University of Portsmouth. School of Pharmacy and Biomedical Sciences, Portsmouth PO1 2DT, UK

## Abstract

The differentiation of oligodendroglia from oligodendrocyte precursor cells (OPCs) to complex and extensive myelinating oligodendrocytes (OLs) is a multistep process that involves largescale morphological changes with significant strain on the cytoskeleton. While key chromatin and transcriptional regulators of differentiation have been identified, their target genes responsible for the morphological changes occurring during OL myelination are still largely unknown. Here, we show that the regulator of focal adhesion, Tensin3 (Tns3), is a direct target gene of Olig2, Chd7, and Chd8, transcriptional regulators of OL differentiation. Tns3 is transiently upregulated and localized to cell processes of immature OLs, together with integrin-β1, a key mediator of survival at this transient stage. Constitutive *Tns3* loss-of-function leads to reduced viability in mouse and humans, with surviving knockout mice still expressing Tns3 in oligodendroglia. Acute deletion of *Tns3 in vivo*, either in postnatal neural stem cells (NSCs) or in OPCs, leads to a two-fold reduction in OL numbers. We find that the transient upregulation of Tns3 is required to protect differentiating OPCs and immature OLs from cell death by preventing the upregulation of p53, a key regulator of apoptosis. Altogether, our findings reveal a specific time window during which transcriptional upregulation of Tns3 in immature OLs is required for OL differentiation likely by mediating integrin-β1 survival signaling to the actin cytoskeleton as OL undergo the large morphological changes required for their terminal differentiation.

## INTRODUCTION

Oligodendrocyte lineage cells, mainly constituted by oligodendrocyte precursor cells (OPCs) and oligodendrocytes (OLs), play key roles during brain development and neuronal support, by allowing saltatory conduction of myelinated axons and metabolically supporting these axons with lactate or glucose shuttling through the myelin sheath (Funfschilling et al., 2012; Lee et al., 2012; Meyer et al., 2018). Accumulating evidence also indicates their fundamental contribution to different aspects of adaptive myelination, a type of brain plasticity (Mount and Monje, 2017; Yang et al., 2020), shown by the requirement of new oligodendrogenesis for proper learning and memory in motor, spatial, and fear-conditioning learning paradigms (McKenzie et al., 2014; Xiao et al., 2016; Steadman et al., 2019; Pan et al., 2020; Wang et al., 2020; Xin and Chan, 2020). Furthermore, oligodendroglia and myelin pathologies have been recently linked, not only to the development of glioma (Liu et al., 2011), but to developmental (Castelijns et al., 2020; Phan et al., 2020), neurodegenerative (Grubman et al., 2019; Bryois et al., 2020), and psychiatric (Nott et al., 2019) diseases.

Unlike most precursor cells, OPCs constitute a stable population of the postnatal and adult CNS (Ffrench-Constant and Raff, 1986; Suzuki and Goldman, 2003). Therefore, OPCs need to keep a tight balance between proliferation, survival, and differentiation. This balance is crucial to maintain the OPC pool while contributing to myelin plasticity in adult life, and to remyelination in diseases such as multiple sclerosis (MS). Demyelinated MS plaques can be normally repaired in early stages of the disease, presumably by endogenous OPCs, but this repair process becomes increasingly inefficient with aging, when OPC differentiation seems to be partially impaired (Chang et al., 2002; Compston and Coles, 2002; Neumann et al., 2019). Therefore, understanding the mechanisms involved in OPC differentiation is critical to foster successful (re)myelination in myelin pathologies.

A large diversity of extrinsic signals, including those mediated by integrin signaling (reviewed in (Bergles and Richardson, 2016)) as well as many intrinsic factors, including transcription factors (TFs) and chromatin remodelers (reviewed in (Emery and Lu, 2015; Parras et al., 2020)), are involved in OPC proliferation, survival, and differentiation. However, the mechanisms for how these signals balance OPC behavior is largely unknown. OPC differentiation requires profound changes in chromatin and gene expression (Emery and Lu, 2015; Küspert and Wegner, 2016; Wheeler and Fuss, 2016). TFs, such as Olig2, Sox10, Nkx2.2 or Ascl1, are key regulators of OL differentiation by directly controlling the transcription of genes implicated in this process (Qi et al., 2001; Stolt et al., 2002; Nakatani et al., 2013; Yu et al., 2013) but being already expressed at the OPC stage, it is still unclear how these TFs control the induction of differentiation. A growing body of evidence suggests that some of these TFs work together with chromatin remodeling factors during transcriptional initiation/elongation to drive robust transcription (Zaret and Mango, 2016). Accordingly, Olig2 and Sox10 TFs have been shown to cooperate with chromatin remodelers such as Brg1 (Yu et al., 2013), Chd7 (He et al., 2016; Marie et al., 2018), Chd8 (Marie et al., 2018; Zhao et al., 2018), and EP400 (Elsesser et al., 2019), to promote the expression of OL differentiation genes. To improve our understanding of the mechanisms of OL differentiation, we searched for novel common targets of these key regulators, by generating and analyzing the common binding profiles of Olig2, Chd7, and Chd8, in gene regulatory elements of differentiating oligodendroglia. We identified *Tns3*, coding for the focal adhesion protein Tensin3, as one such target and showed that it is expressed in immature OLs during myelination and remyelination, thus constituting a marker for this transient oligodendroglial stage. Using different genetic strategies to induce *Tns3* loss-of-function mutations *in vivo*, we describe, the function of a Tensin family member in the CNS, demonstrating that Tns3 is required for oligodendrocyte differentiation in the postnatal mouse brain, at least in part by mediating integrin-β1 signaling, essential for survival of differentiating oligodendroglia (Colognato et al., 2002; Benninger et al., 2006).

## RESULTS

### *Tns3* is a direct target gene of key regulators of oligodendrocyte differentiation

To find new factors involved in oligodendrocyte differentiation, we screened for target genes of Olig2, Chd7, and Chd8, key regulators of oligodendrogenesis (Lu et al., 2000; Lu et al., 2002; Yu et al., 2013; He et al., 2016; Marie et al., 2018; Zhao et al., 2018; Parras et al., 2020). We generated and compared the genome-wide binding profiles for these factors in acutely purified oligodendroglial cells from postnatal mouse brain cortices by magnetic cell sorting (MACS) of O4^+^ cells (Marie et al., 2018). MACS-purified cells, composed of 80% PDGFRα^+^ OPCs and 20% of Nkx2.2^+^/CC1^+^ immature oligodendrocytes (iOLs), were subjected to chromatin immunoprecipitation followed by sequencing (ChIP-seq) for Olig2 and histone modifications marking the transcription activity of gene regulatory elements (H3K4me3, H3K4me1, H3K27me3, and H3K27ac; Fig. 1A). The profile of activity histone marks at Olig2 binding sites indicated that Olig2 binds promoters (60%) and enhancers (40%) with either active or more poised/repressive states (Fig. S1A-F), supporting the suggested pioneer function of Olig2 in oligodendrogenesis (Yu et al., 2013). Among the 16,578 chromatin sites bound by Olig2 corresponding to 8,672 genes (Fig. S1D), there were key regulators of oligodendrocyte differentiation, including *Ascl1, Sox10, Myrf, Chd8*, and *Smarca4/Brg1* (Fig. 1B; supplementary table 1). Combining Olig2 with Chd7 and Chd8 binding profiles, that we previously generated using the same protocol (Marie et al., 2018), we found 1774 chromatin sites commonly bound by the three regulators, with half of them (47% and 832 peaks) corresponding to active promoters (H3K4me3/H3K27ac marks) of 654 protein-coding genes (Fig. 1C, supplementary table 2). Among these genes, *Tns3* coding for Tensin3, a focal adhesion protein deregulated in certain cancers (Martuszewska et al., 2009), showed the highest expression levels in iOLs relative to other brain cell types (Zhang et al., 2014; Fig. 1D). Indeed, Olig2, Chd7, and Chd8 commonly bound three putative promoters of *Tns3* having active transcription marks in purified oligodendroglia (H3K27ac/H3K24me3; Fig. 1E). To directly assess whether *Tns3* expression requires the activity of these key regulators, we interrogated the transcriptomes of these oligodendroglial cells purified from *Chd7iKO* (*Pdgfra-CreER^T^*; *Chd7^flox/flox^*), *Chd8cKO (Olig1^Cre^; Chd8 ^flox/flox^*), and their respective control cortices (Marie et al., 2018; Zhao et al., 2018). *Tns3* transcripts were largely downregulated upon acute deletion of these factors in postnatal OPCs/iOLs (Fig. 1F,G), indicating that *Tns3* expression in OPCs/iOLs is directly controlled by Chd7 and Chd8 chromatin remodelers, key regulators of oligodendrocyte differentiation.

**Figure 1.**
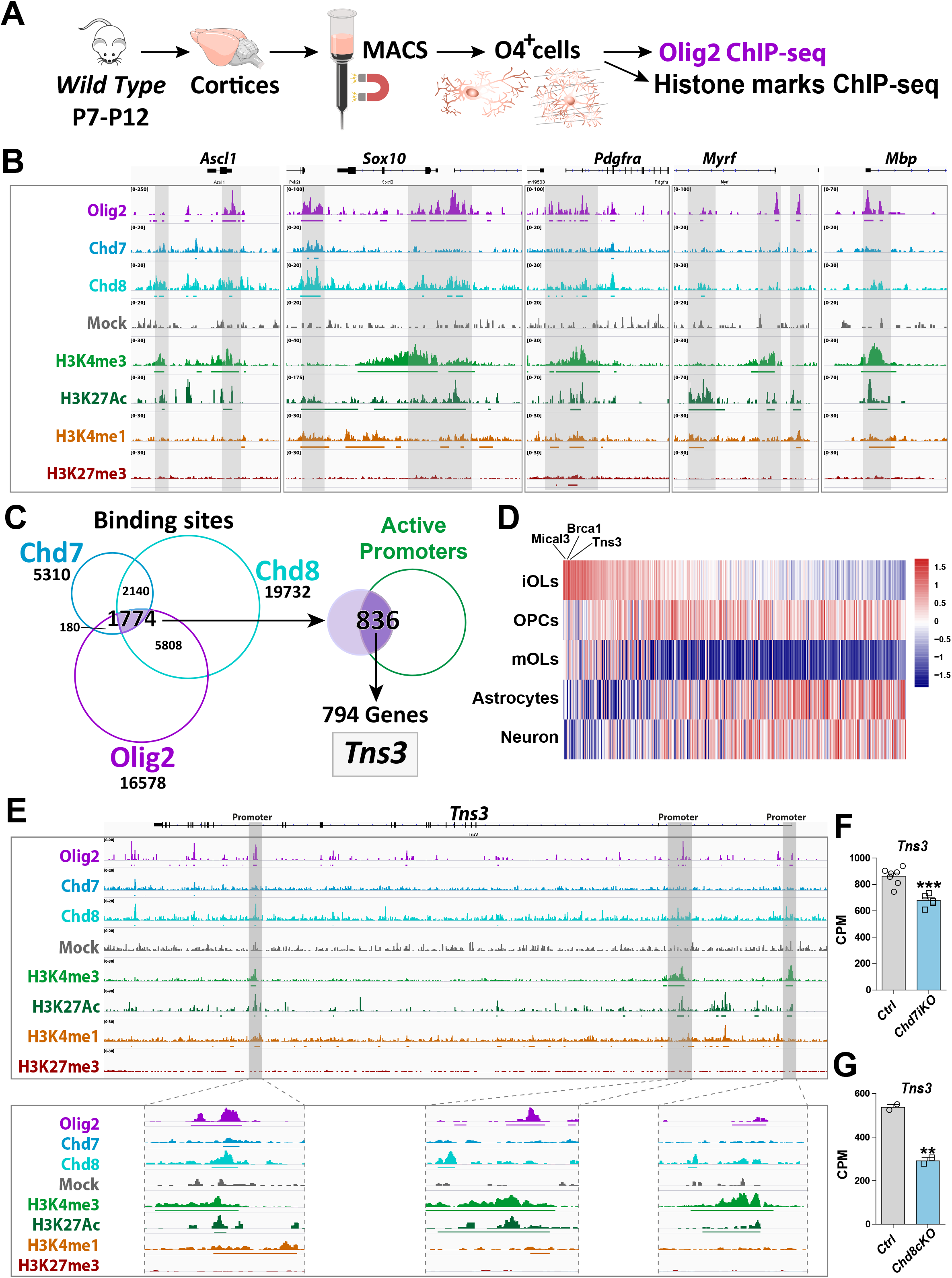
*Tns3* is a target gene of Olig2 and Chd7/8 regulators of oligodendrocyte differentiation. **(A)** Scheme representing MACSorting of O4^+^ cells from wild type cortices followed by ChIP-seq for Olig2 and histone marks (H3K4me3, H3K27Ac, H3K4me1 and H3K27me3). **(B)** Tracks from IGV browser of *Ascl1, Sox10, Pdgfra, Myrf and Mbp* gene regions depicting ChIP-seq data in O4^+^ cells (OPCs/OLs) for transcription factor Olig2, chromatin remodelers Chd7 and Chd8 and epigenetic marks (H3K4me3, H3K27ac, H3K4me1 and H3K27me3). Mock (control IgG) shows no peaks in the regions of interest. Lines present below peaks indicate statistical significance (by peak calling). **(C)** Strategy used to identify *Tns3* as a gene target of Olig2, Chd7, and Chd8, potentially involved in oligodendrogenesis. Left, Venn diagrams depicting the overlap of binding peaks between Chd7 (blue), Chd8 (cyan), and Olig2 (purple) in O4^+^ cells. Right, Venn diagram showing that 836 (47%) of the 1774 common regions have marks of active promoters, corresponding to 794 genes, including *Tns3*. **(D)** Heatmap representing the expression of the 794 genes in iOLs compared to OPCs, mOLs, astrocytes and neurons. *Tns3* is the third most specific (data from Zhang et al., 2014). **(E)** Tracks from IGV browser of *Tns3* gene region depicturing ChIP-seq data in O4^+^ cells (OPCs/OLs) for transcription factor Olig2 and epigenetic marks (H3K4me3, H3K27ac, H3K4me1, and H3K27me3), zooming in Tns3 alternative promoters. Mock (control IgG) shows no peaks in the regions of interest. Horizontal lines present below peaks indicate statistical significance (peak calling). **(F,G)** Barplots showing *Tns3* transcript count per million (CPM) in O4^+^ cells upon tamoxifen-induced *Chd7* deletion (*Chd7iKO*, (**F**) or *Chd8* deletion (*Chd8cKO*, (**G**) compared to control (Ctrl). Statistics done using EdgeR suite.

### *Tns3* transcripts are highly expressed in mouse and human immature oligodendrocytes

We then investigated *Tns3* expression pattern in the brain. High expression levels of *Tns3* transcripts in iOLs, compared to its low expression in other cells of the postnatal mouse brain detected by bulk transcriptomics (Fig S2A; Zhang et al., 2014) were paralleled by the sparse labelling with *Tns3* probes enriched in the white matter of the postnatal and adult brain detected by *in situ* hybridization (Fig. S2B; Allen Brain atlas, https://portal.brain-map.org/. By harnessing single-cell transcriptomics (scRNA-seq), we sought to create an integrative gene profiling for oligodendrocyte lineage cells, by bioinformatics integration and analyses of oligodendrocyte lineage cells at embryonic, postnatal, and adult stages (Marques et al., 2016, 2018). We thus integrated these datasets using Seurat (Stuart et al., 2019) and selected 5,516 progenitor and oligodendroglial cells. Unsupervised clustering and visualization of cells in two dimensions with uniform manifold approximation and projection (UMAP), identified nine different clusters following a differentiation trajectory. Based on known cell-subtype specific markers (Fig. S2E and supplementary Table 3), we could identify these clusters as (Fig. S2C,E): (1) two types of neural stem/progenitor cells, that we named NSCs and NPCs according to their expression of stem cell (*Vim, Hes1, Id1*) and neural progenitor (*Sox11, Sox4, Dcx*) markers; (2) OPCs expressing their known markers (*Pdgfra, Cspg4, Ascl1*) and cycling OPCs also enriched in cell cycle markers (*Mki67, Pcna, Top2)*; (3) two stages of iOLs, both expressing the recently proposed markers *Itpr2* and *Enpp6* (Marques et al., 2016; Xiao et al., 2016), and which are split by the expression of *Nkx2-2* (iOL1 being *Nkx2-2*^+^ and iOL2 being *Nkx2-2*^-^), in agreement with our previous characterization by immunofluorescence (Nakatani et al., 2013; Marie et al., 2018); (4) myelin forming oligodendrocytes (MFOLs), enriched in markers such as *Slc9a3r2* and *Igsf8*; and (5) two clusters of myelinating OLs, that we named MOL1 and MOL2, expressing transcripts of myelin proteins (*Cnp, Mag, Mbp*, *Plp1*, *Mog*) and some specific markers of each cluster, including *Mgst3, Pmp22* for MOL1 (corresponding to MOL1/2/3/4 clusters of Marques 2016), and *Neat1*, *Grm3*, *Il33* for MOL2 (corresponding to MOL5/6 clusters of Marques 2016). Interestingly, *Tns3* transcripts were strongly expressed in both iOL1 and iOL2 clusters (Fig. S2D,E), similar to the recently proposed iOL markers *Itpr2* and *Enpp6* (Fig. S2E), and downregulated in mature/terminally differentiated oligodendrocytes, indicating that high levels of *Tns3* expression are specific to iOLs. We finally, assessed whether *Tns3* expression pattern was conserved in human oligodendroglia pursuing a similar bioinformatics analysis using single cell transcriptomes from human oligodendroglia differentiated from embryonic stem cells (Chamling et al., 2021). Upon integration with Seurat and identification of cluster cell types using specific markers, we selected 7,690 progenitor and oligodendroglial cells that corresponded to six main cluster cell types following a differentiation trajectory from neural cells (NSCs) up to immature OLs (iOL1 and iOL2), as depicted by UMAP representation (Fig. S2F,H). Cells expressing high levels of *TNS3* corresponded to immature oligodendrocytes (iOL1 and iOL2 clusters, Fig. S2G,H). We obtained similar results analyzing a human fetal midterm cerebellum (GW9-GW22) dataset (Aldinger et al., 2021), with high levels of *TNS3* in iOLs similar to other suggested iOL markers such as *ITPR2*, *ENPP6*, and *BCAS1*, indicating a conserved expression pattern of Tns3/TNS3 between mouse and human oligodendrogenesis.

### Tns3 protein is enriched in the cytoplasm and processes of immature oligodendrocytes

Given the high expression level of *Tns3* transcripts in immature oligodendrocytes, we characterized the Tns3 protein expression pattern in the postnatal brain using commercial and homemade Tns3-recognizing antibodies. Optimization of immunofluorescence protocols demonstrated signal in CC1^+^ oligodendrocytes in the postnatal brain with four different antibodies (P24, Fig. S3). To our surprise, while all antibodies showed signal localized in the cytoplasm and main processes of CC1+ OLs (Fig. S3A-E), one Tns3-recognizing antibody (Millipore) also presented a strong nuclear signal never reported for Tns3 localization in other tissues (such as lung, liver, and intestine) (Katz et al., 2007; Nishino et al., 2012; Cao et al., 2015). To better characterize Tns3 protein expression pattern and its subcellular localization, we generated a knock-in mice tagging the Tns3 C-terminal side with a V5-tag (*Tns3^Tns3^*^-V5^ mice) by microinjecting mouse zygotes with a single strand oligodeoxynucleotide (ssODN) containing V5 sequence together with Cas9 protein and a gRNA targeting the stop codon region of *Tns3* (Methods; Fig. S4A-C). We first verified by immunofluorescence that Tns3-V5 protein in *Tns3^Tns3^*^-V5^ mice presented the expression pattern reported for Tns3 in the lung and the kidney (Fig. S4D,E). We then characterized Tns3 protein expression in oligodendroglia using V5 antibodies, finding that Tns3 protein can be detected at high levels in the cytoplasm and main processes of CC1^+^ iOLs but not in their nuclei (Fig. 2A). Using an antibody recognizing Itpr2, a suggested iOL marker (Marques et al., 2016), we saw that Tns3 largely overlapped with Itpr2 (Fig. 2B). Using Nkx2.2 and Olig1^cytoplamic^ expression distinguishing iOL1 and iOL2 respectively, we found high levels of Tns3 in iOL1s (Nkx2.2^+^/Olig1^-^ cells) and a fraction of iOL2s (Nkx2.2^-^/Olig1^cytoplamic^ cells; Fig. 2C), suggesting that Tns3 protein expression peaks in early iOLs. Comparison with Opalin protein localized in the cell body, processes, and myelin segments of oligodendrocytes, showed that Tns3 levels decreased with increasing levels of Opalin, with Tns3-V5 levels undetectable in myelinating oligodendrocytes (i.e. Opalin^+^/CC1^+^ cells presenting myelinated segments; Fig. 2D, arrowheads). We then performed Western blot analysis with anti-V5 antibodies in purified O4^+^ cells from P7, P14, and P21 *Tns3^Tns3^*^-V5^ mouse brains to assess their specificity to recognize Tns3-V5, knowing that two Tns3 isoforms can be detected at the transcript level in the human brain (Fig. S4F, GTEX project, gtexportal.org/home/gene/TNS3). Indeed, we could detect both the full-length (1450 aa, 155 kDa) and the Tns3 short (C-term, 61 kDa) isoforms in O4+ cells from brains at P7 and P14 stages having many iOLs, but not at P21 having mainly mOLs (Fig. S4G,H), thus validating the specificity of the anti-V5 antibodies in recognizing Tns3 protein. We eventually found a Tns3 antibody also recognizing the C-terminal of Tns3 protein (Sigma Ct) that upon optimized immunofluorescence labeling confirmed the Tns3 expression pattern seen with the V5 antibodies. In combination with Nkx2.2 and Olig1 immunofluorescence, it showed that Tns3 is strongly detected in the cytoplasm and main cellular processes of all iOL1s, defined as Nkx2.2^high^/Olig1^-^ cells having a round nucleus and small cytoplasm (Fig. 2E, white arrows), and it divided iOL2s, defined as Nkx2.2^-^/CC1^high^ cells, into three stages: (1) Tns3^high^/Nkx2.2^-^/Olig1^-^ (Fig. 2E, arrowheads), (2) Tns3^high^/Nkx2.2^-^/Olig1^high-cytoplamic^ (Fig. 2E, grey arrows), and (3) Tns3^-^ /Nkx2.2^-^/ Olig1^high-cytoplamic^ (Fig. 2E). A similar Tns3 expression pattern and localization was found *in vitro* using neonatal neural progenitors’ differentiating cultures, where Tns3 was detected together with CNP myelin protein in the cytoplasm and cell processes of Nkx2.2^high^/CNP^+^ differentiating oligodendrocytes (Fig. 2F,G). Altogether, these results indicate that high but transient levels of Tns3 protein characterize early immature oligodendrocytes (iOL1s and early iOL2s), being localized at their cytoplasm and cell processes (Fig. 2H,I).

**Figure 2.**
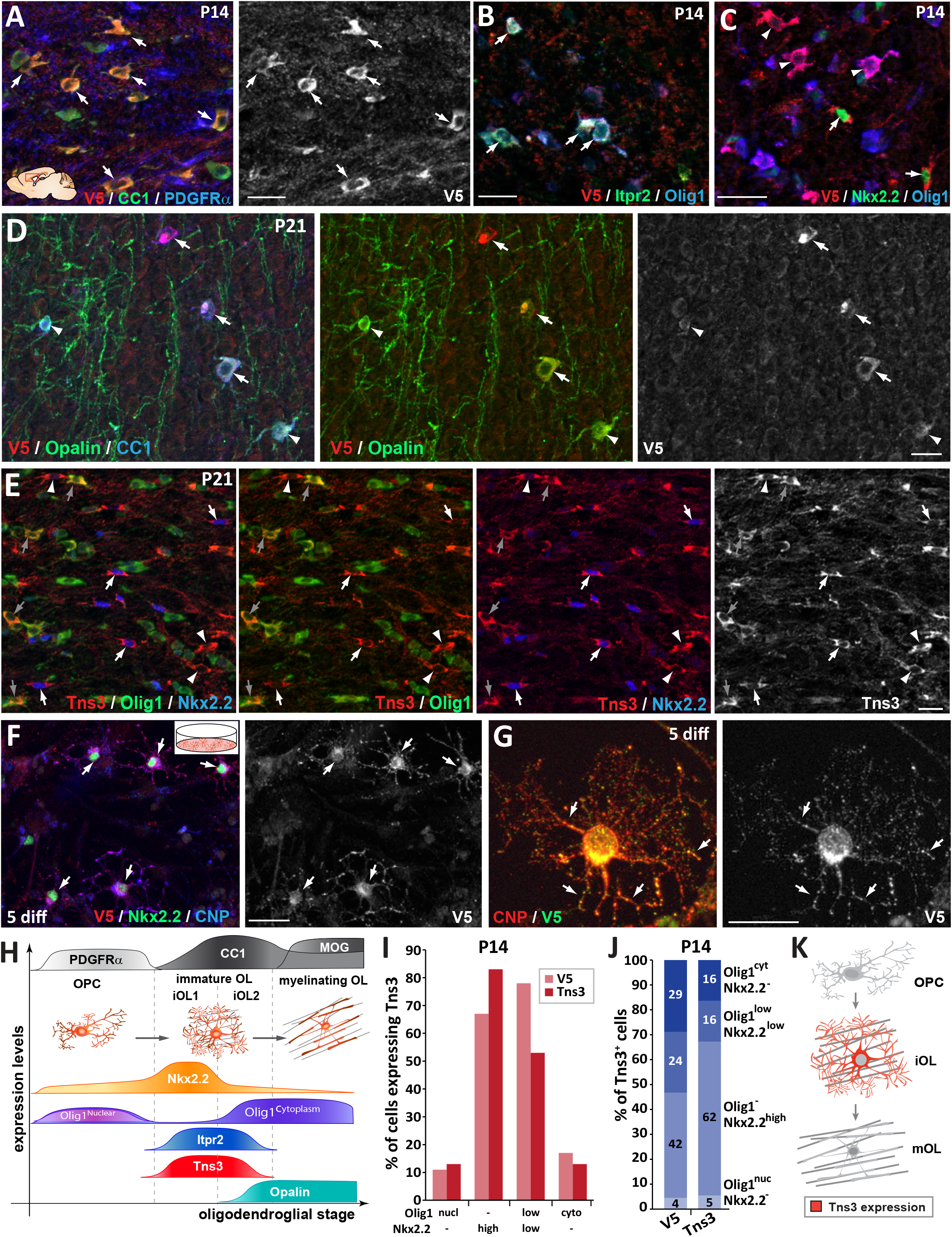
Tns3 protein is detected at high levels in the cytoplasm and main cell processes of immature oligodendrocytes in the postnatal brain. Immunofluorescence in sagittal sections of postnatal brain at the level of the corpus callosum at P14 **(A-C)** and P21 **(D-E)** using V5 and Tns3 antibodies. **(A)** Tns3-V5 is detected at high levels in CC1^+^ OLs (arrows) but not in PDGFRα^+^ OPCs. **(B)** Tns3-V5 expression overlap well with Itpr2 (arrows), with some of them being Olig1^high-cytoplamic^ cells. **(C)** Tns3-V5 overlaps with high levels of Nkx2.2 expression (arrows) and also with Nkx2.2^-^/Olig1^high-cytoplamic^ cells (arrowheads). **(D)** Tns3-V5 expression overlap with Opalin in iOLs (arrows, CC1^+^ cells with large cytoplasm) but is downregulated in Opalin^+^ mOLs (arrowheads, CC1^+^ cells with small cytoplasm and myelin segments). **(E)** Tns3 (Sigma-Ct antibody) is detected at high levels in Nkx2.2^+^/Olig1^-^ early iOL1s (white arrows), in late Nkx2.2^-^/Olig1^-^ iOL1s (white arrowheads), and in Nkx2.2^-^/Olig1^high-cytoplamic^ iOL2s (grey arrows). **(F)** Tns3-V5 expression NSC cultures after 5 days in differentiation. Note the Tns3 expression in Nkx2.2^+^/CNP^+^ OLs (arrows). **(G)** Subcellular localization of Tns3 expression in CNP^+^ OLs present in the cytoplasm and in dots distributed along the cell processes, overlapping with CNP signal (arrows). **(H)** Schematic representation of Tns3 expression together with key markers of different oligodendroglial stages summarizing data shown in A-E panels. **(I)** Histogram representing the percentage of Nkx2.2^-^/Olig1^high-nuclear^, Nkx2.2^high^/Olig1^-^, Nkx2.2^low^/Olig1^low-cytoplamic^ and Nkx2.2^-^/Olig1^high-cytoplamic^ cells expressing Tns3-V5 and Tns3 at P14. **(J)** Histogram representing the percentage of Tns3-V5^+^ and Tns3^+^ cells at P14 that are Nkx2.2^-^/Olig1^high-nuclear^, Nkx2.2^high^/Olig1^-^, Nkx2.2^low^/Olig1^low-cytoplamic^ and Nkx2.2^-^/Olig1^high-cytoplamic^. **(K)** Schematic representation of Tns3 expression and subcellular localization in oligodendroglia. Scale bars: A-F, 20 μm; G, 10 μm.

Finally, we investigated whether other Tensin family members were expressed in oligodendroglia, finding that Tns1 and Tns2 but not Tns4 were detectable at low levels in iOLs by immunofluorescence (Fig. S5A,B), paralleling their low transcription levels compared to *Tns3* (Fig. S5C; brainrnaseq.org). Therefore, Tns3 appears to be the main Tensin expressed during oligodendrocyte differentiation, suggesting that Tns3 function in immature oligodendrocytes is likely to be evolutionarily selected, and thus of biological importance in oligodendrogenesis.

### Tns3 expression is found in immature oligodendrocytes during remyelination

Given the strong Tns3 expression in iOLs during postnatal myelination, we hypothesized that Tns3 expression could be enriched during remyelination in newly formed oligodendrocytes contributing to remyelination. To test this hypothesis, we performed lysolecithin (LPC) focal demyelinating lesions in the corpus callosum of adult (P90) *Tns3^Tns3^*^-V5^ and wild-type mice, and assessed for Tns3 expression at the peak of oligodendrocyte differentiation (8 days post-lesion) in this remyelinating model (Nait-Oumesmar et al., 1999). We found that while non-lesioned adult brain regions contained only sparse Tns3^+^ iOLs (CC1^high^/Olig1^cyto-high^ cells), remarkably many Tns3^+^ iOLs were detected in the remyelinating area using both V5 (Fig. S6A,C, arrows) and Tns3 antibodies (Fig. S6B, arrows). Quantification of Tns3^+^ cells showed a clear increase in Tns3^+^ iOLs around the lesion borders compared to the corpus callosum far from the lesion area (Fig. S6D), suggesting that Tns3 expression may be a useful marker of on-going remyelination and lesion repair. Of note, we could also detect Tns3 expression in some microglia/macrophages in the lesion area using a combination of F4/80 antibodies (Fig. S6C, arrowheads). Altogether, all these data indicate that Tns3 expression peaks at the onset of oligodendrocyte differentiation, labeling immature oligodendrocytes during both myelination and remyelination.

### *In vivo* CRISPR-mediated *Tns3* loss-of-function in neonatal neural stem cells impairs oligodendrocyte differentiation

To explore the role of Tns3 in OL differentiation, we first utilized a *Tns3* gene trap mouse line (*Tns3^βgeo^*; Chiang et al., 2005), and two CRISPR-mediated indel mutation mice presumptively leading to Tns3 constitutive knockout. Analyses of these three mouse lines (Fig. S7; see Methods for details) showed both developmental lethality (in line with loss-of-function variants of TNS3 causing ∼80% developmental mortality in the human population; https://gnomad.broadinstitute.org; Fig. S8; Methods) and possible genetic compensation in Tns3 expression, making them inappropriate tools to study Tns3 function in oligodendrogenesis.

Given the tendency of cells to escape the *Tns3* loss-of-function upon constitutive knockout mutations, we decided to assess *Tns3* requirement during postnatal oligodendrogenesis by inducing *in vivo* acute *Tns3*-deletion in few neural stem cells (NSCs) of the neonatal brain and tracing their cell progeny with a GFP reporter. For this, we combined the postnatal electroporation technique with CRISPR/Cas9 technology. First, we used our previously validated gRNAs targeting *Tns3* at the first coding ATG (exon 6; Fig. S9; Methods) inserting them in an integrative CRISPR/Cas9 plasmid also expressing the GFP reporter (Fig. 3A), to transfect neonatal NSCs of the dorsal subventricular zone (SVZ), which generate a large number of oligodendroglial cells during the first postnatal weeks (Kessaris et al., 2006; Nakatani et al., 2013), and focused our study on glial cells by quantifying the GFP^+^ progeny of targeted NSCs, outside the SVZ and located in the dorsal telencephalon three weeks later (P22, Fig. 3B). The fate of GFP^+^ cells was determined by immunodetection of GFP and glial subtype markers (CC1^high^ for oligodendrocytes, PDGFRα for OPCs, and CC1^low^ and their unique branched morphology for astrocytes). Remarkably, brains electroporated with the CRISPR plasmids targeting *Tns3* had a 2-fold reduction in GFP^+^ oligodendrocytes compared to brains electroporated with control plasmids, while GFP^+^ OPCs were found in similar proportions (Fig. 3C,C’,D). The proportion of GFP^+^ astrocytes was increased by 1.5-fold, likely as a result of the large reduction in oligodendrocytes, as the number of GFP^+^ astrocytes was not changed (61.3 ± 10.9 in experimental versus 57.2 ± 11.8 in controls; Fig. 3C,C’,D). To assess whether the reduction in oligodendrocytes from *Tns3*-deleted NSCs was the consequence of a reduction in OPCs generated, we assessed for possible changes in numbers, proliferation, and survival of OPC at P11, when most cortical OPCs have not yet started differentiation. We found no differences in the proportion of GFP cells being OPCs (Fig. 3E), nor the proliferative status of GFP^+^ OPCs (MCM2^+^/PDGFRα^+^ cells; Fig. 3F) between experimental and control brains, while the reduction of oligodendrocytes was already marked (Fig. 3E), indicating that loss of *Tns3* only affected the process of OPC differentiation into oligodendrocytes.

**Figure 3.**
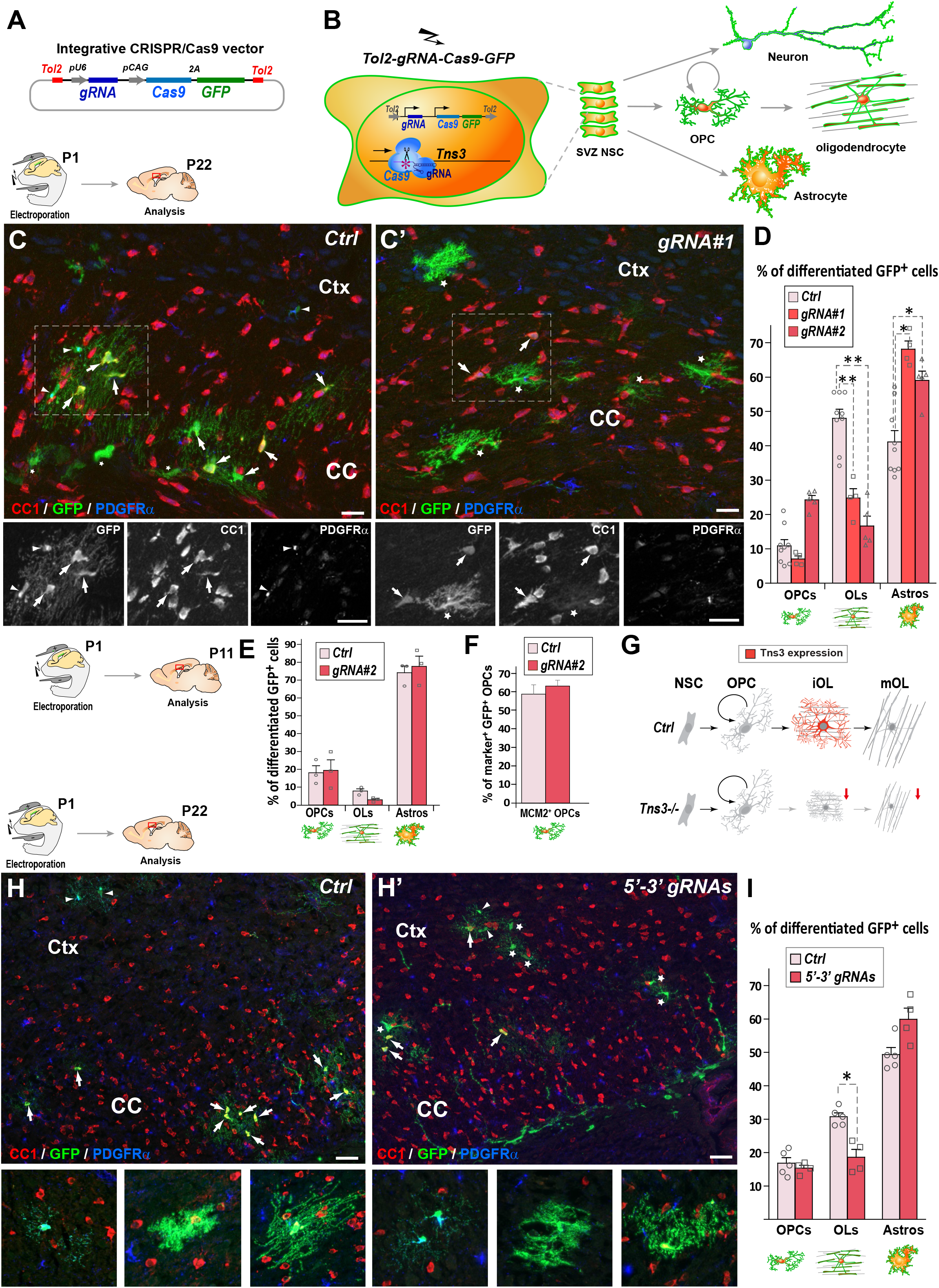
CRISPR-mediated *Tns3* mutation in NSCs reduces oligodendrocyte differentiation in the postnatal brain. **(A)** Schematic of the CRISPR/Cas9 expression vector allowing Tol2-DNA integration driving Cas9 and GFP expression (from polycistronic 2A-mediated cleaved) from CAG promoter and sgRNA expression from U6 promoter. **(B)** Schematic of the dorsal SVZ NSC electroporation of CRISPR-plasmid at postnatal day 1 (P1) and traced neural cell-subtype progeny. **(C,C’)** Immunofluorescence of representative P22 sagittal sections of the dorsal telencephalon showing GFP^+^ cells being either PDGFRα^+^ OPCs (arrowheads), CC1^high^ OLs (arrows) or CC1^low^ astrocytes (asterisks) progeny of P1 NSCs electroporated either with *Ctrl* plasmid **(C)** or *Tns3-gRNA#1* plasmid **(C’)**. **(D)** Histogram showing the percentage of GFP^+^ glial cell types found in *Ctrl*, *gRNA#1* or *gRNA#2* electroporated brains being PDGFRα^+^-OPCs, CC1^high^-OLs and CC1^low^-astrocytes. Note the 2-fold reduction of CC1^+^ OLs in *Tns3-gRNA* transfected brains, as illustrated in C’ compared to C. **(E)** Histograms representing the percentage of GFP^+^ differentiated cells at P11. Note the lack of changes in OPCs, and the incipient reduction in OLs. **(F)** Histograms quantifying the proportion of proliferative (MCM2^+^ cells) GFP^+^ OPCs in electroporated P11 mice brain. **(G)** Schematic of Tns3 expression in mice (Upper) and of the effects of *Tns3* CRISPR-mediated deletion (Lower). **(H,H’)** Representative P22 sagittal sections of the dorsal telencephalon showing GFP^+^ cells being either PDGFRα^+^ OPCs (arrowheads), CC1^high^ OLs (arrows) or CC1^low^ astrocytes (asterisks) progeny of P1 NSCs electroporated either with *Ctrl* plasmid **(H)** or *Tns3-5’-3’* targeting plasmid (labeled as *5’-3’ gRNAs*) **(H’)**. **(I)** Histograms showing the percentage of GFP^+^ glial cell types found in *Ctrl* or 3’- 5’ *gRNA* electroporated brains being PDGFRα^+^-OPCs, CC1^high^ OLs and CC1^low^ astrocytes in the corpus callosum (CC) and cortex (Ctx). Note the 2-fold reduction of CC1^+^ OLs in *Tns3-5’-3’ gRNA* transfected brains. Scale bar, 20 μm.

Given the expression of two Tns3 isoforms in the brain (Fig. S3F,G), we asked whether a deletion of both isoforms would have a greater impact in oligodendrocyte differentiation. We thus used two gRNAs efficiently cutting the beginning and the end of *Tns3* coding sequence (5’-3’gRNAs, Methods), to delete the whole *Tns3* locus. We found a similar reduction of oligodendrocytes in the loss of the two *Tns3* isoforms than in mutations affecting only full-length Tns3 (Fig. 3H,H’,I), suggesting that the small Tns3 isoform does not play an additional role with full-length Tns3 in oligodendrocyte formation. Altogether, these results indicate that *Tns3* loss-of-function mutations in neonatal SVZ-NSCs impair OPC differentiation without apparent changes in OPC generation and proliferation, thus suggesting that Tns3 is largely required for OPC differentiation into oligodendrocytes in the postnatal brain (Fig. 3G).

### OPC-specific *Tns3* deletion impairs oligodendrocyte differentiation in the postnatal brain

Given the heterogeneity of CRISPR/Cas9-mediated indels and the difficulties to assess *in vivo* the penetrance of their *Tns3* loss-of-function, to address in more detail the consequences of *Tns3* loss-of-function, we designed a *Tns3* conditional knockout allele, by flanking with LoxP sites the exon 9 (Fig. S10). In order to specifically delete *Tns3* in postnatal OPCs, we administered tamoxifen at P7 to *Pdgfra-CreER^T^; Tns3^flox/flox^; Rosa26^stop^*^-*YFP*^ (hereafter called *Tns3-iKO* mice) and control pups (*Pdgfra-CreER^T^; Tns3^+/+^; Rosa26^stop^*^-*YFP*^ littermates) and analyzed its effects on oligodendrogenesis at P14 and P21 (Fig. 4A) both in white matter (corpus callosum and fimbria) and grey matter regions (cortex and striatum). We first assessed for the efficiency of *Tns3* deletion in Nkx2.2^+^/GFP^+^ iOLs from different regions by immunofluorescence using a Tns3 antibody (Sigma Ct), finding that the strong Tns3 signal present in Nkx2.2^+^/GFP^+^ iOLs of control brains was almost completely eliminated in *Tns3-iKO* iOLs without affecting Tns3 expression in vessels (Fig. S11B,B’,C, arrows and arrowheads versus asterisks). We then assessed for changes in oligodendrogenesis. Remarkably, the number of oligodendrocytes (CC1^+^/GFP^+^ cells) was reduced by half in all quantified regions (reduction of 38.95% in the CC, 48.60% in cortex, 50.88% in the fimbria, 38% in the striatum; Fig. 4B-D, Fig. S11E-F’, Fig. S12B,B’) in *Tns3-iKO* compared to control, while OPC (PDGFRα^+^/GFP^+^ cells) density was unchanged (Fig. 4B-D). Using markers distinguishing different stages of oligodendrocyte differentiation (iOL1 and iOL2/mOL), we found that the density of iOL1s (Nkx2.2^high^ cells), which express the highest levels of Tns3 protein in control brains, was unchanged (Fig. S11B,B’,D), while the density of early iOL2s (CC1^+^/Olig1^-^ cells) and later oligodendrocyte stages (iOL2/mOLs CC1^+^/Olig1^+^ cells), was reduced by 30% and 50% respectively in *Tns3-iKO* compared to controls (Fig. 4E-F), suggesting that Tns3 is required for normal oligodendrocyte differentiation. Finally, we assessed possible changes in OPC proliferation by immunodetection of Mcm2, finding no significant changes in the proliferation of *Tns3-iKO* OPCs compared to control OPCs (Fig. S12C,C’,D).

**Figure 4.**
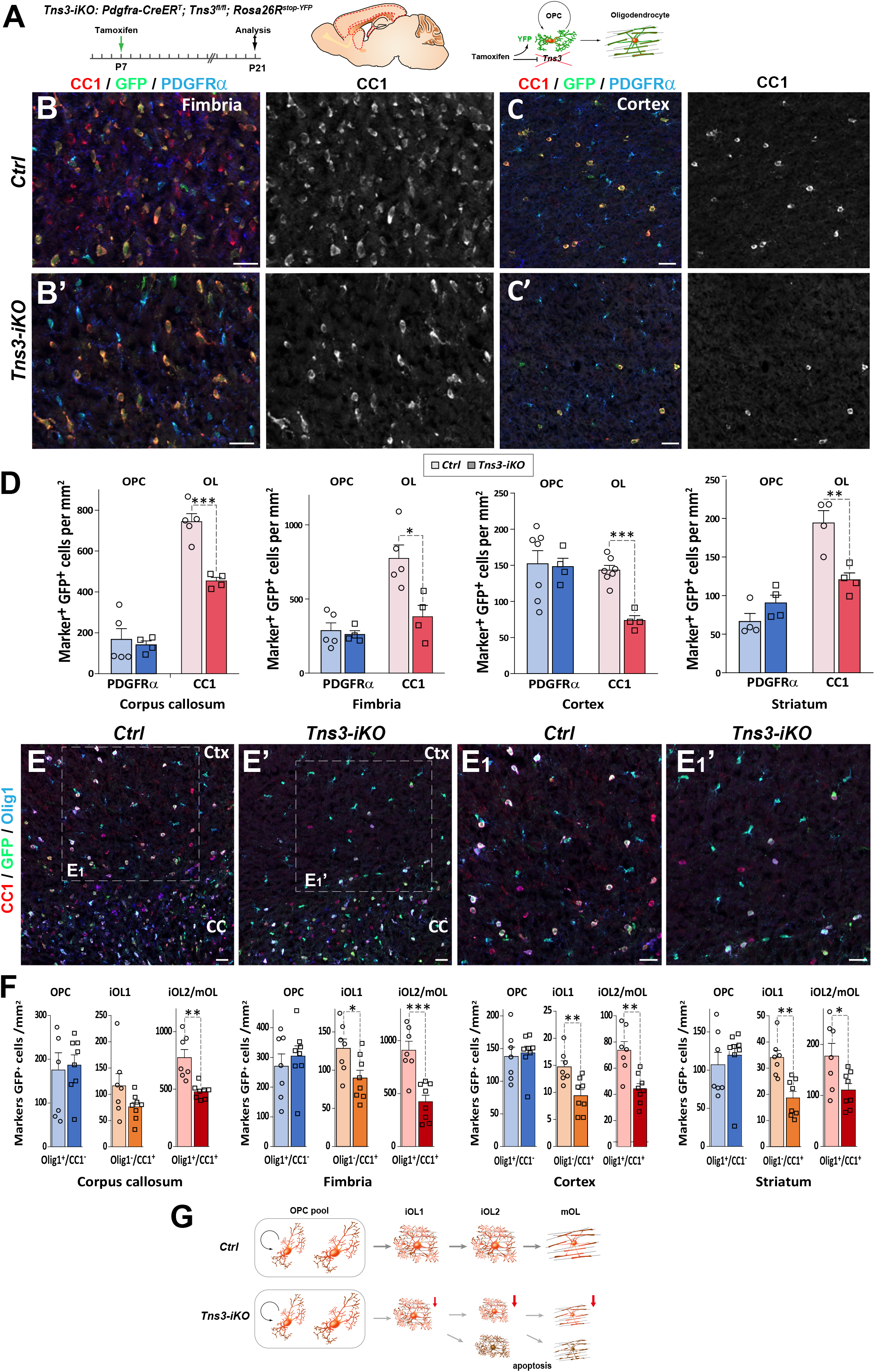
OPC-specific *Tns3* deletion reduces the number of differentiating oligodendrocytes in the postnatal brain. **(A)** Scheme of tamoxifen administration to *Tns3-iKO* and control (*Cre^+^; Tns3^+/+^*) mice, Cre-mediated genetic changes, and timing of experimental analysis. **(B-B’, C-C’)** Immunofluorescence in P21 sagittal brain sections for CC1, GFP and PDGFRα illustrating similar density of OPCs (PDGFRα^+^) and 2-fold reduction in OL (CC1^+^) density in *Tns3-iKO* **(B’,C’)** compared to control **(B,C)** in the fimbria **(B)** and the cortex **(C)**. **(D)** Histograms showing OPC and OL density in P21 *Tns3-iKO* and control (*Ctrl*) mice, in the corpus callosum, fimbria, cortex, and striatum. Note the systematic OL decrease of 40-50% in each region. **(E-E_1_’)** Immunofluorescence in P21 sagittal brain sections for Olig1, GFP and CC1 to distinguish three stages of oligodendrogenesis: OPCs (Olig1^+^/CC1^-^), iOL1s (CC1^+^/Olig1^-^) and iOL2s/mOLs (CC1^+^/Olig1^+^) in *Ctrl* **(E)** or *Tns3-iKO* mice **(E’)**. E_1_ and E_1_’ are higher magnification of the squared area in E and E’. **(F)** Histograms showing the OPCs, the iOL1s and the iOL2s/mOLs density in P21 *Tns3-iKO* and control mice, in the corpus callosum, fimbria, cortex, and striatum. Note the decrease of iOL1s and iOL2s over 40% in each area quantified (except for iOL1 density in the corpus callosum). **(G)** Schematic representing defects in oligodendrogenesis found in *Tns3-iKO* compared to control. CC, corpus callosum; Ctx, cortex. Scale bar, 20 μm.

Altogether, these results indicate that acute deletion of *Tns3* in OPCs reduces by 2-fold generation of oligodendrocytes in the postnatal brain, without major changes in OPC numbers and proliferation (Fig. 4G).

### *Tns3-iKO* oligodendroglia undergo apoptosis

Tensins are known to mediate integrin stabilization and activation in other cells-types (Liao and Lo, 2021), with Tns3 been shown to bind integrin-β1 through its phosphotyrosine-binding domain and FAK through its SH2 domain in fibroblasts (Cui et al., 2004; Liao et al., 2007; Georgiadou et al., 2017). In oligodendroglia, integrin α6β1 association with Fyn kinase is required to amplify PDGF survival signaling and to promote myelin membrane formation, by switching neuregulin signaling from a PI3K to a MAPK pathway (Colognato et al., 2004). Moreover, by conditional ablation of integrin-β1 *in vivo*, it was demonstrated that integrin-β1 signaling is involved in survival of differentiating oligodendroglia, but not required for axon ensheathment and myelination *per se* (Benninger et al., 2006). We therefore, investigated the expression of genes involved in integrin signaling in the transcriptome of oligodendroglial cells. Indeed, *Tns3* expression pattern in iOLs was closely matching that of *Itgb1* (integrin-b1), *Fyn*, *Bcar1*/*p130Cas*, and *Ptk2*/*Fak* both in mouse and human oligodendroglia (Fig. S13A,B). Furthermore, using neural progenitor differentiation cultures, we observed co-expression of integrin-β1 and Tns3 in CNP^+^ oligodendrocytes by immunofluorescence (Fig. S13C), suggesting that Tns3 could relay integrin-β1-mediated survival signal in differentiating oligodendroglia. Therefore, we assessed for signs of cell death in *Tns3-iKO* oligodendroglia by performing the TUNEL technique together with GFP and CC1 immunodetection. Interestingly we found a 5-fold increase in TUNEL^+^ cells in the dorsal telencephalon of *Tns3-iKO* brain, compared to control, without significant changes in non-oligodendroglial cells present in the SVZ (Fig. 5A-C). To gain more insight into the cellular alterations and cell death of *Tns3-*deleted oligodendroglia, we investigated their cellular morphology and behavior by video microscopy during their differentiation in culture. To this end, we MACS-purified OPCs from *Tns3-iKO* and control (*Tns3^flox/flox^; Rosa26^stop^*^-YFP^ littermates) mice at P7, two days after administration of tamoxifen, plated them in proliferating medium for three days, and recorded their behavior during three days in the presence of differentiation medium (Fig. 5D). Using the expression of the YFP as a readout of Cre-mediated recombination, we compared the behavior of YFP^+^ cells (*Tns3-iKO*) with neighboring YFP^-^ cells (internal control) in the same cultures. In parallel, we used MACSorted cells from control mice as external control. Quantification of the proportion of YFP^+^ cells over time showed a 20% reduction of YFP^+^ cells (from 80% to 60%) during the 3 days in proliferation medium followed by a reduction to 50% by day 3 in differentiation medium (Fig. 5E), suggesting possible cell death of *Tns3*-mutant cells. Live imaging monitoring of cell behavior showed that once YFP^+^ cells had developed multiple branched morphology, characteristic of differentiating oligodendrocytes, they showed a 4-fold increase in their probability to die compared to YFP^−^ cells of the same culture (Fig. 5F-H, yellow and white arrows, respectively) or to cells from control cultures, with more pronounced cell death by the third day of culture (Fig. 5F,G). Together, these results indicate that *Tns3-iKO* oligodendroglia present increased cell death both *in vivo* and in primary culture, at the stage when Tns3 is upregulated and cells start to developed their branched morphology, suggesting that Tns3 likely mediates β1-integrin signaling required for their survival.

**Figure 5.**
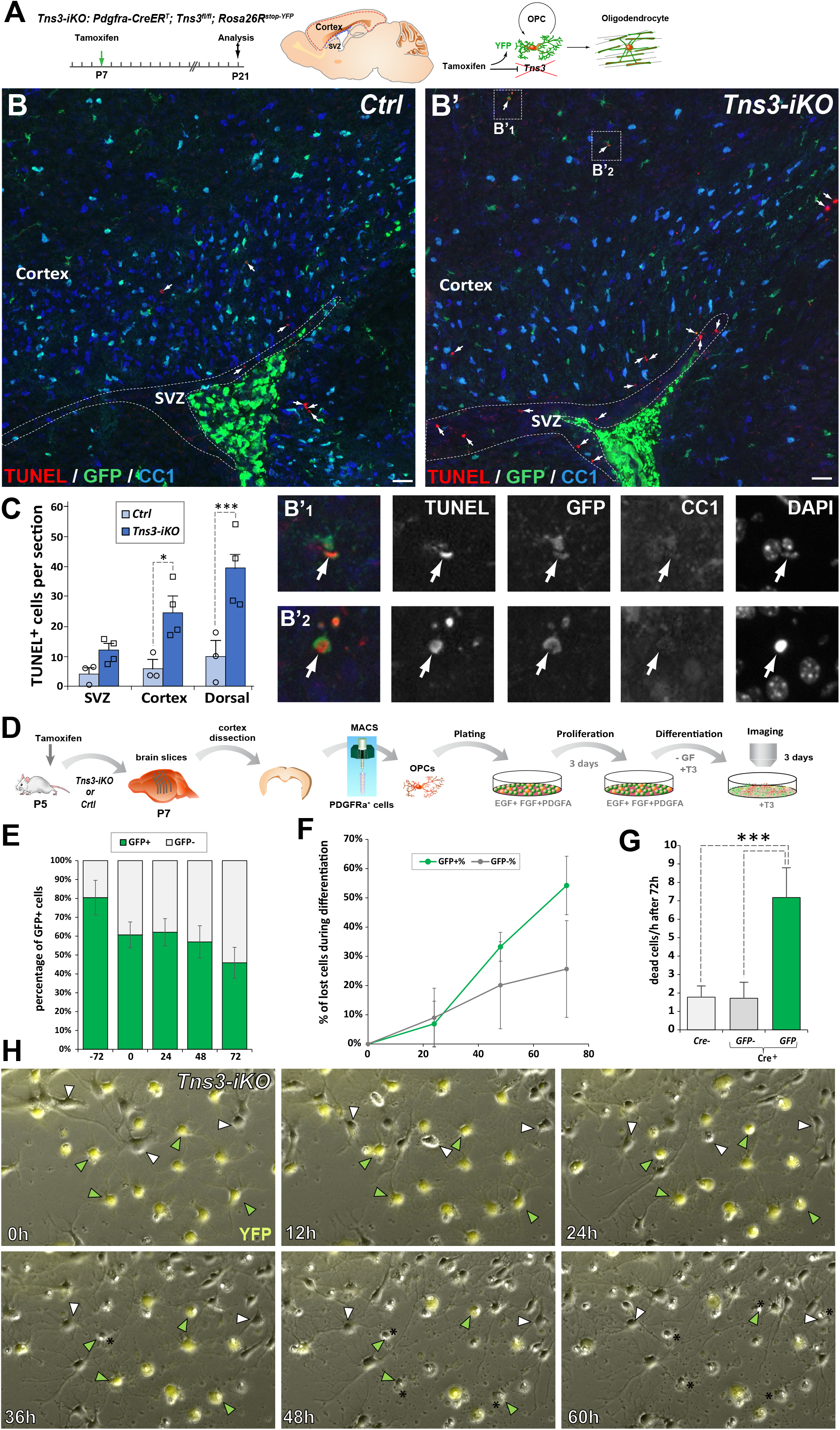
Increased cell death of *Tns3-iKO* oligodendroglia. **(A)** Scheme of tamoxifen administration to *Tns3-iKO* (*Cre^+^; Tns3^fl/fl^*) and control (*Cre^+^; Tns3^+/+^*) mice, Cre-mediated genetic changes, and timing of experimental analysis. **(B-B’)** Immunodetection of death cells by TUNEL, together with recombined cells (GFP) and OLs (CC1^+^ cells) showing increase number of TUNEL^+^ cells in the cortex and corpus callosum of *Tns3-iKO* mice **(B’)** compared to control mice **(B)**, including some GFP^+^/CC1^−^ OPCs (insets B’_1_ and B’_2_). **(C)** Histograms quantifying the number of TUNEL^+^ cells in the SVZ, cortex and both together (dorsal) per section. **(D)** Scheme of the video microscopy protocol in MACSorted OPCs purified from *Tns3-iKO* and control P7 mice. Cells were imaged every 10 minutes during 72h in differentiation medium. **(E)** Histograms showing the reduction of the GFP^+^ cells proportion from the plating (−72h) to the end of the experiment (72h after differentiation onset). **(F)** Time curve quantifying the loss of GFP^+^ OLs compared to GFP^-^ OLs during the 72h of differentiation. **(G)** Histograms representing the quantification of cells lost per hour during the 72h differentiation period, showing a 5-fold increase in loss of GFP^+^ *Tns3-iKO* cells compared to GFP^-^ cells (non-recombined cells from *Tns3-iKO* mice, internal negative control) or cells coming from *Cre^−^* littermates (*Cre^−^*, external negative control). **(H)** Time lapse frames showing cells every 12 hours illustrating both GFP^+^ (green arrowheads, recombined *Tns3-iKO* cells) and GFP^-^ (white arrowheads, non-recombined *Tns3-iKO* cells) that die over the time of video microscopy. Note the larger number of GFP^+^ OLs (cells with multi-branched OL morphology) dying compared to GFP^-^ OLs. SVZ, subventricular zone. Scale bar, 20 μm.

### Apoptosis of *Tns3-iKO* oligodendroglia is mediated by p53 upregulation

To study the molecular mechanisms of Tns3 function in oligodendroglia, we first looked at P53 expression, the master transcriptional regulator of the cellular genotoxic stress response (Kastenhuber and Lowe, 2017; Aubrey et al., 2018). Interestingly, we found a 10-fold increase in p53^+^ OPCs (GFP^+^/CC1^-^ cells) and 4-fold increase in p53^+^ iOLs (GFP^+^/CC1^+^ cells) in *Tns3-iKO* compared to control (Fig. 6A-C), suggesting that the loss of Tns3 leads to an upregulation of p53, that together with the loss of integrin-β1 survival signal, mediates the cell death of *Tns3-iKO* differentiating oligodendroglia (Fig. 6D).

**Figure 6.**
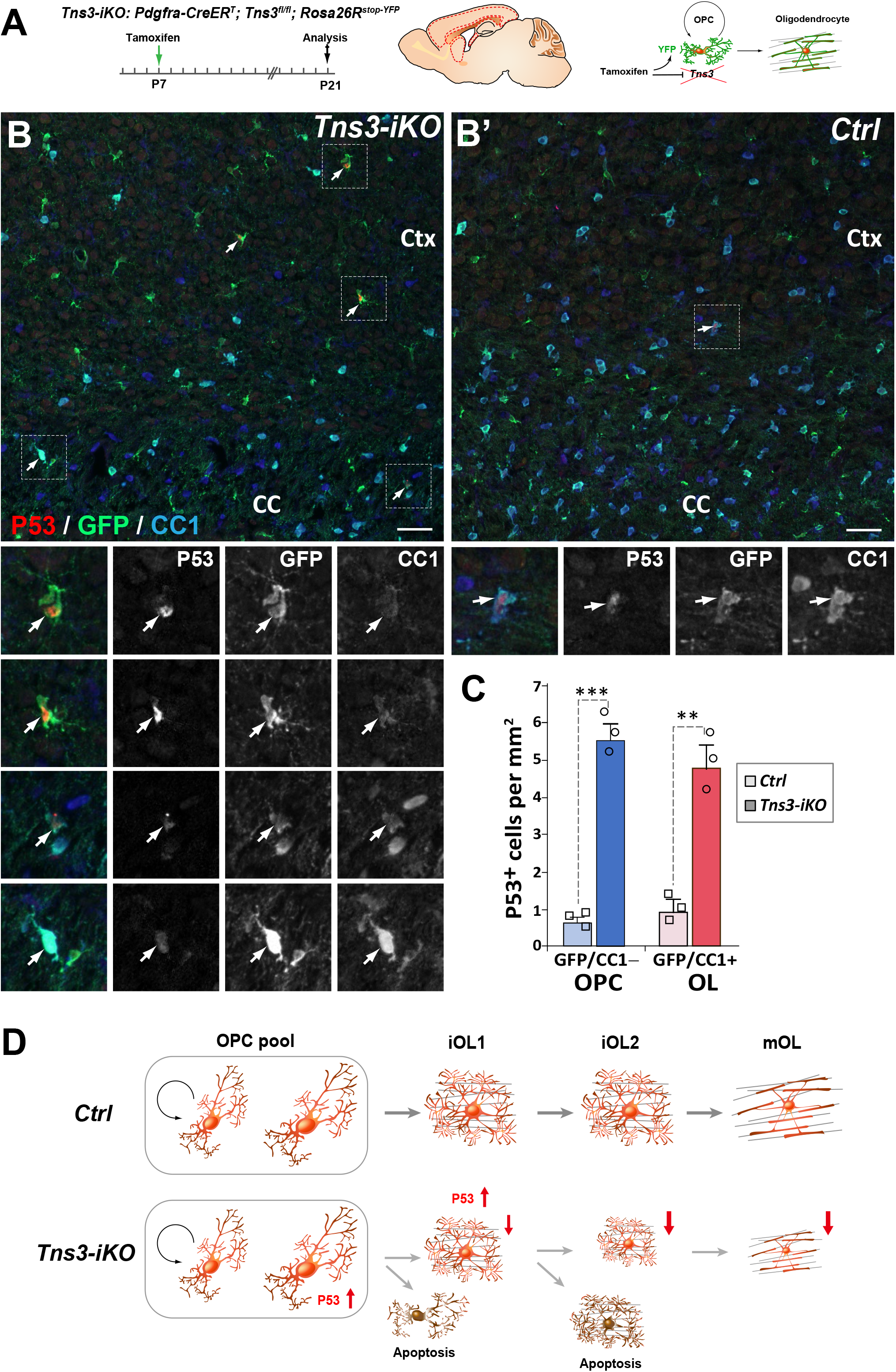
p53-mediated cell death of *Tns3-iKO* oligodendroglia. **(A)** Scheme of tamoxifen administration to *Tns3-iKO* (*Cre^+^; Tns3^fl/fl^*) and control (*Cre^+^; Tns3^+/+^*) mice, Cre-mediated genetic changes, and timing of experimental analysis. **(B-B’)** Immunodetection of p53, together with GFP to label recombined cells and CC1 to label OLs showing a strong increase number of p53^+^/GFP^+^/CC1^−^ OPCs and p53^+^/GFP^+^/CC1^+^ OLs in the cortex and corpus callosum of *Tns3-iKO* mice **(B)** compared to control mice **(B’)**. Dotted squares highlight some cases of p53^+^ cells, shown at higher magnification below. **(C)** Histograms quantifying the number of p53^+^ cells per area (mm^2^) in the dorsal telencephalon. CC, corpus callosum; Ctx, cortex. Scale bar, 20 μm. **(D)** Schematics representing Tns3-deleted phenotypes in oligodendroglia. CC, corpus callosum; Ctx, cortex. Scale bar, 20 μm.

### *Tns3-iKO* oligodendroglia shows transcriptional dysregulation of genes involved in OPC differentiation, apoptosis, integrin signaling, and cell cycle regulation

To further unravel the defects of *Tns3*-deleted oligodendroglia, we purified oligodendroglia (O4^+^ cells) from P12 *Tns3-iKO* and control cortices by magnetic cell sorting (MACS) (Fig. 7A). Upon validation of Tns3 deletion at the transcript and protein levels (Fig. S15A-C) and that similar proportion of oligodendroglia were present in each genotype (Fig. S15D-F), we compared their transcriptomes obtained by bulk RNA sequencing (Fig. 7A). Principal component analysis of Tns3-iKO and control samples show clear separation between the groups (Fig. S15G) and statistical analyses using EdgeR (Chen et al., 2016) showed and 2082 differentially expressed genes (DEGs, p-value < 0.05) between *Tns3-iKO* and control, with 834 downregulated and 1248 upregulated genes (Fig. 7B). Gene ontology analysis of biological processes indicated that main downregulated processes were involved in terms related OL differentiation (including gliogenesis, glial cell differentiation, oligodendrocyte differentiation, lipid metabolism, and positive regulation of cell projection organization; Fig. 7C,E, supplemental table 4), while the upregulated biological processes related to terms such as cellular stress and p53 pathway (including double strand break repair, cellular response to oxidative stress, and signal transduction of p53 class mediator), opposite processes involved in cell cycle regulation (including DNA integrity check point, G2/M transition of mitotic cell cycle, and positive- and negative-regulation of cell cycle process) (Fig. 7D,F,G, supplemental table 5). Interestingly, while GO processes related to integrin signaling and cell adhesion were upregulated (including positive regulation of cell adhesion, regulation of cell adhesion mediated by integrin, integrin-mediated signaling pathway, and positive regulation of cell adhesion; Fig. 7D, supplemental table 5, integrin sheet), with several integrin transcripts upregulated (*Itgam, Itga8, Itgb2, Itga4, Itgb3, Itgb5, Itga6, Itgb8*, which are normally not expressed in OPCs/iOLs; supplemental table 5, integrin sheet graphic), while *Fyn*, Src family kinase that associates with α6β1 and is required to amplify PDGF survival signaling (Collognato et al., 2004), was downregulated (1.4-fold, p-value = 0.03; supplemental table 5, integrin sheet), suggesting that Tns3 deletion impairs normal integrin signaling, and as a consequence Tns3-deleted cells try to compensate this impairment upregulating of other integrin family members. These results, confirming and expanding those obtained by immunofluorescence analyses, led us to propose a model suggesting that *Tns3*-deleted oligodendroglia present signs of cellular stress accompanied by double strand break signaling upregulation (including ATM and CHK2 regulators), p53 stabilization and upregulation of p53 target genes involved in apoptosis (including PUMA, APAF1, and Caspase 7), and conflicting signals related to cell cycle (upregulation of p21, promoting cell cycle arrest, and upregulation of CDK/Cyclin complexes promoting cell cycle progression).

**Figure 7.**
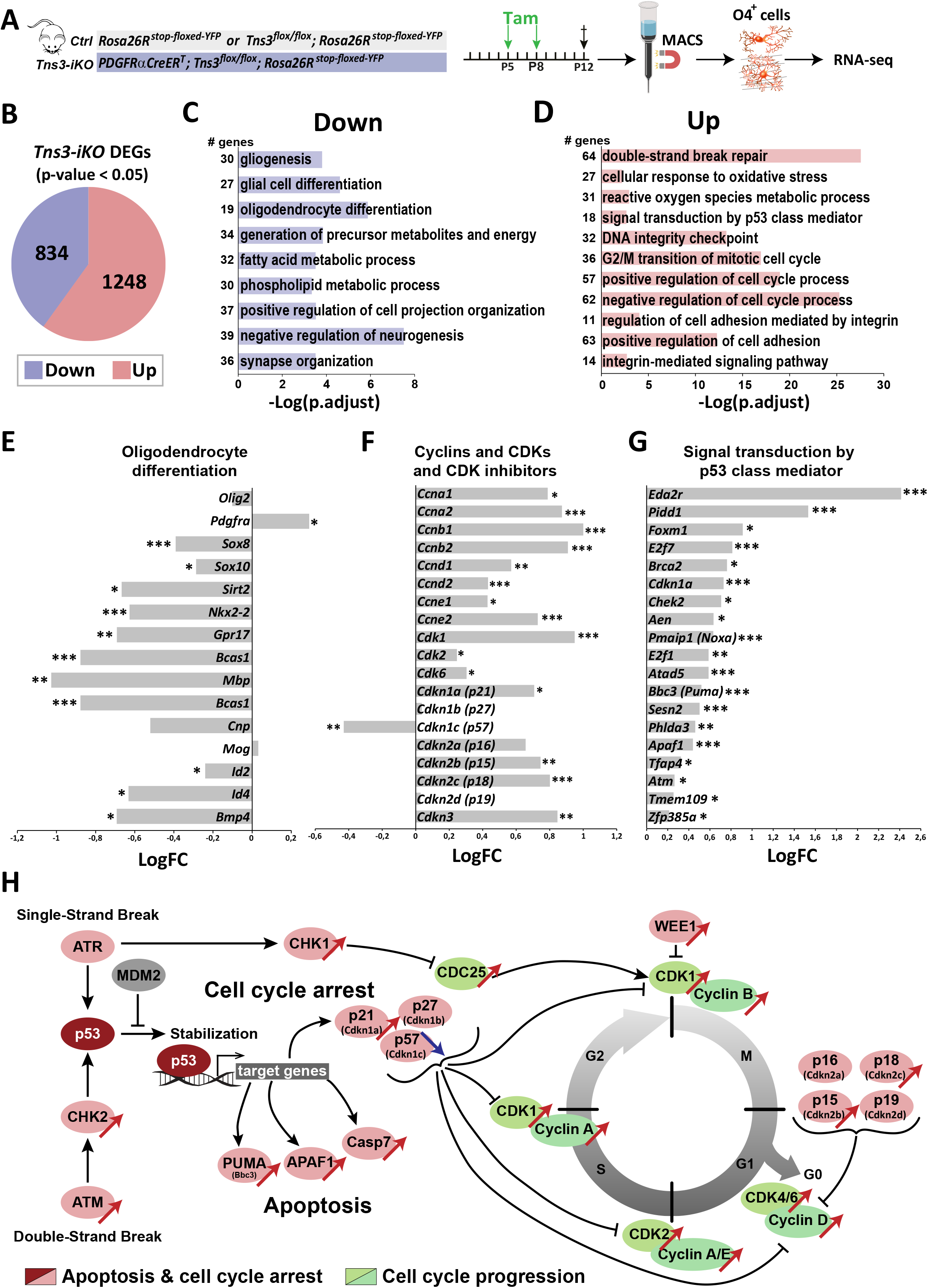
Mechanisms involved in *Tns3-iKO* oligodendroglial defects. **(A)** Diagram representing tamoxifen (Tam) injection in P5 and P8 *Ctrl* and *Tns3-iKO* pups followed by MACSorting of O4^+^ cells which were used to do RNA-seq. **(B)** Pie chart showing the amount of differentially expressed genes (DEGs; p-value < 0.05) in *Tns3-iKO* O4^+^ cells compared to *Ctrl*. **(C-D)** Gene ontology (GO) analysis of downregulated (**C**) and upregulated (**D**) biological processes in *Tns3-iKO* O4^+^ cells compared to *Ctrl*. Numbers on left represent the number of deregulated genes in each GO process. **(E-G)** Graphs representing the logarithmic fold change (LogFC) for an example of genes involved in oligodendrocyte differentiation (E), cell cycle (F) and p53 pathway (G). **(H)** Summary schematics of the transcriptional dysregulation of *Tns3-iKO* O4^+^ cells, representing the upregulation of the apoptosis pathway and the conflicting signals on cell cycle arrest/progression. Red and blue arrows represent gene upregulation and downregulation respectively, in *Tns3-iKO* O4^+^ cells compared to *Ctrl*.

## DISCUSSION

The tight balance of oligodendrocyte precursor cells (OPCs) between proliferation, survival, and differentiation ensures their capacity to respond to the myelination needs of the CNS by generating new oligodendrocytes on demand, whilst avoiding the generation of brain gliomas through uncontrolled OPC proliferation. The observation that OPCs are present within demyelinating MS lesions, but fail to efficiently differentiate into myelinating cells with age and disease progression (Chang et al., 2002; Neumann et al., 2019), together with the strong sensitivity of immature oligodendrocytes to survival/apoptotic signals (Hughes and Stockton, 2021), suggests that efforts to foster OPC differentiation and survival of immature oligodendrocytes are a critical events for healthy aging and successful remyelination in MS patients. In this study, we combined the genome-wide binding profile of key regulators of oligodendrocyte differentiation, Olig2, Chd7, and Chd8 (Lu et al., 2000; Zhou et al., 2000; Lu et al., 2002; Zhou and Anderson, 2002; He et al., 2016; Küspert and Wegner, 2016; Marie et al., 2018; Zhao et al., 2018), to identify their common gene targets, and focused our analysis on Tensin3 (Tns3), whose expression matched the onset of oligodendrocyte differentiation. To study Tns3 expression and function, we generated several genetic tools, including CRISPR/Cas9 vectors to induce *Tns3* mutations both *in vivo* and *in vitro*, a *Tns3^Tns3^*^-V5^ knock-in mouse, two constitutive *Tns3* knockout mice, and finally an inducible knockout (*Tns3^Flox^*) mouse. Using these tools, we provide several lines of evidence showing that Tns3 is upregulated in immature oligodendrocytes (iOLs) and required for normal oligodendrocyte differentiation. First, we show that Tns3 expression is strongly induced at the onset of oligodendrocyte differentiation, localized to the cytoplasm and main cell processes of iOLs, and downregulated in mature oligodendrocytes both at the transcript and protein levels, thus constituting a novel marker for iOLs, for which we provide an optimal immunofluorescence protocol with a commercial antibody (Sigma, Ct). Second, we show that during remyelination, Tns3 is also expressed in newly formed oligodendrocytes and thus could be used as a hallmark for ongoing remyelination. Third, analyzing both *Tns3^βgeo^* gene trap mice and two *Tns3^KO^* mice, we show that constitutive *Tns3* deletion is detrimental for normal development and that the predicted loss of Tns3 full-length transcript and protein is bypassed in the oligodendroglia of surviving homozygous animals, paralleling the intolerance for *TNS3* loss-of-function variants found in the human population. Fourth, *in vivo* CRISPR-mediated *Tns3*-deletion in neonatal neural stem cells from the subventricular zones leads to a 2-fold reduction of oligodendrocytes without changes in OPC generation, proliferation, and numbers. Fifth, *in vivo Tns3* induced knockout (*Tns3-iKO*) in postnatal OPCs leads to a 2-fold reduction of differentiating oligodendrocytes without reducing the overall OPC population, both in grey and white matter brain regions. Finally, we provide evidence, by immunodetection *in vivo* and video microscopy of primary OPC differentiation cultures, that *Tns3-iKO* differentiating oligodendroglia upregulate p53, key sensor of cell stress, and present a 4-5-fold increase in apoptosis compared to control oligodendroglia, suggesting that mechanistically Tns3 function is likely required for normal oligodendrocyte differentiation at least in part by mediating integrin-β1 survival signaling in differentiating oligodendroglia.

### Tns3 is a novel marker for immature oligodendrocytes

Recent studies has started to uncover genes enriched in immature oligodendrocytes (iOLs), such as *Itpr2* (Zeisel et al., 2015; Marques et al., 2016), *Enpp6* (Xiao et al., 2016), and *Bcas1* (Fard et al., 2017), that could be used as markers for these transient cell populations, particularly interesting to label areas of active (re)myelination in the context of oligodendrocyte and myelin pathology, such as preterm brain injury and multiple sclerosis. Here, we report for the first time that Tns3 is a hallmark of iOLs (figure 2). Tns3 is expressed at high levels in iOLs and downregulated as oligodendrocytes mature into myelinating cells, showing a complete overlap with Itpr2 transcript and protein. We found that a commercial Tns3 antibody (Millipore) also recognizes another nuclear protein that, like Tns3 in the cytoplasm, also labels at high levels iOLs, paralleling the case of CC1 antibody, which recognizes both APC and Quaking-7 proteins in oligodendrocytes (Lang et al., 2013; Bin et al., 2016). Upon testing several antibodies, we found one (Sigma Ct) optimally labeling iOLs by immunofluorescence in brain sections and oligodendroglial cell cultures, whereas the Itpr2 commercial antibody we tried did not match this high quality iOLs immunolabeling. An optimized protocol for immunodetection using Bcas1-recognizing antibodies has been shown to label iOLs (Fard et al., 2017). Finally, Enpp6 is very specific for iOLs at the transcript level (Xiao et al., 2016) but to our knowledge, no Enpp6-recognizing antibodies producing good quality immunodetection are yet available. Therefore, Tns3 protein expression in the CNS is a hallmark of iOLs, and the Tns3 Sigma Ct antibody is an optimal reagent to label iOLs during both myelination and remyelination.

### Tns3 is required for oligodendroglial differentiation

Oligodendrocyte differentiation involves substantial generation of new membrane and cell processes composing the 40-60 myelin segments formed by mature oligodendrocytes (Hughes et al., 2018). Actin cytoskeleton remodeling is an important driver of the OL morphological changes undergone during their differentiation (Nawaz et al., 2015; Zuchero et al., 2015). Tensin proteins, linking the extracellular signals received by transmembrane integrins with the actin cytoskeleton in different cell types (Liao and Lo, 2021), are well placed to play an important role in these morphological changes. At the molecular level, it has been shown that the phosphotyrosine-binding domains of tensins interact with the NPXY motifs present in the cytoplasmic tails of integrin-β1 in a pTyr-insensitive fashion (Calderwood et al., 2003; Katz et al., 2007; McCleverty et al., 2007), allowing tensins to bring actin filaments, through their actin binding domain, to focal adhesion sites (Liao and Lo, 2021). Given that the extension of OL cell processes’ growth cone is guided by the sequential activation of Fyn, FAK and RhoGAP (Thomason et al., 2020), and that high levels of Tns3 protein are detected in the cell body and processes of iOLs coinciding with their large enlargement, Tns3 is thus well placed to mediate integrin signaling to the actin cytoskeleton and play an active role in this large cellular remodeling. Moreover, Integrin-β1, FAK/Ptk2, Fyn, p130Cas/Bcar1, and Tns3 are all highly expressed in iOLs (Fig. S13). Here, using three independent approaches, we show that loss of Tns3 in iOLs reduces by half the numbers of oligodendrocytes in the postnatal brain. It is therefore very likely that Tns3 act as a mediator of integrin α6β1 signaling to promote OL survival and differentiation by mediating actin cytoskeletal remodeling. Finally, through its additional ability to bind to EGFR (Cui et al., 2004), whose activation is another driver of oligodendroglial differentiation, Tns3 could also be required for mediation of signaling downstream of growth factor receptor activation in early iOLs; this could also explain the increased death in OLs lacking Tns3.

### Tns3 role in immature oligodendrocyte survival

Programmed cell death regulates developmental oligodendrogenesis, with a large proportion of immature oligodendrocytes degenerating before the fourth week of postnatal life in mice (Barres et al., 1992; Trapp et al., 1997). Also in the adult mouse brain, differentiating OPCs remain in the immature oligodendrocyte stage for roughly 2 days with many of them undergoing programmed cell death (Hughes et al., 2018), indicating that this immature stage is very dependent on survival signals. Apoptotic pathways involving BCL-2 family members have been shown to regulate this oligodendroglial programmed cell death (reviewed in (Hughes and Stockton, 2021). One study has shown that the transcription factor TFEB, involved in autophagy and lysosomal biogenesis, ensures the spatial and temporal specificity of developmental myelination by promoting the expression of ER stress genes and PUMA, a pro-apoptotic factor inducing Bax-Bak-dependent programmed cell death in differentiating oligodendroglia (Sun et al., 2018). Another recent study showed that during both during homeostasis and remyelination, the activity of the primary sensor of cellular stress, nuclear factor (erythroid-derived 2)-like 2 (Nrf2), induces the expression of Gsta4, a scavenger of lipid peroxidation, with in turn controls apoptosis of immature oligodendrocytes via the mitochondria-associated Fas-Casp8-Bid-axis (Carlström et al., 2020).

Several studies have shown that integrin-β1 signaling is required for iOL survival. Neuronal-derived signals, including neuregulin and laminin-2, are received by immature oligodendrocytes through integrin-β1 signaling that would enhance the function of neuroligin as a survival factor by inducing a survival-dependence switch from the phosphatidylinositol 3-kinase-Akt pathway to the mitogen-activated protein kinase (MAPK) pathway, with enhanced MAPK signaling inactivating the pro-apoptotic molecule BAD (Colognato et al., 2002; Benninger et al., 2006). Also, PDGF survival signaling in OPCs and myelin formation have been shown to depend on integrin α6β1 binding to Fyn (Colognato et al, 2004). Tensins typically reside at focal adhesions, which connect the extracellular matrix (ECM) to the cytoskeletal networks through integrins and their associated protein complexes (Kumar, 1998; Liao and Lo, 2021), with focal adhesions mediating both outside-in and inside-out signaling pathways that regulate cellular events, such as cell attachment, migration, proliferation, apoptosis, and differentiation (Liao and Lo, 2021). In this study, we show that Tns3 expression timing during oligodendrogenesis, parallels that of integrin-β1, and we provided evidence of Tns3 and integrin-β1 co-localization in dotted structures resembling nascent and focal adhesion in the cytoplasm and processes of immature oligodendrocytes. Moreover, Tns3-deleted oligodendroglia have a 1.4-fold downregulation of *Fyn* transcripts, Src family kinase which associates with α6β1 and is required to amplify PDGF survival signaling (Collognato et al., 2004), accompanied by the upregulation of transcripts other integrin family member normally not expressed in OPCs and immature OLs (supplemental table 5), suggesting cellular changes to compensate the integrin signaling impairment of *Tns3*-deleted cells. Remarkably, knockout mice for *integrin-α6* present a 50% reduction in brainstem MBP^+^ OLs at E18.5, just before they die at birth, accompanied by an increase in TUNEL^+^ dying OLs (Colognato et al., 2002), while conditional deletion of *integrin-β1* in immature OLs by *Cnp-Cre* also leads to a 50% reduction in cerebellar OLs at P5, with a parallel increase in TUNEL^+^ dying OLs (Benninger et al., 2006). Therefore, given that *Tns3*-induced deletion in postnatal OPCs also leads to 40-50% reduction in OLs in both grey and white matter regions of the postnatal telencephalon (this study), paralleled by similar increase in TUNEL^+^ apoptotic oligodendroglia, we suggest that Tns3 is required for integrin-β1 mediated survival signal in immature oligodendrocytes. Moreover, we suggest that this would lead to cellular stress of *Tns3*-deleted differentiating oligodendroglia and to the upregulation of p53, master regulator of cellular stress and apoptosis, which has been previously been shown to be involved in the apoptosis of human oligodendrocytes in the context of MS (Ladiwala et al., 1999; Wosik et al., 2003) and in the cuprizone demyelination mouse model (Li et al., 2008; Luo et al., 2021).

In summary, here we have generated powerful genetic tools allowing to assess for the first time the role of Tns3 in the CNS, shown that Tns3 protein is found at high levels in the cytoplasm and main processes of immature oligodendrocytes thus constituting a new marker of this oligodendroglial stage, and demonstrated by different genetic approaches that Tns3-deletion leads to a two-fold reduction in differentiating oligodendrocytes, explained at least in part by their increased apoptosis due to p53 upregulation and likely the loss of integrin-β1-mediated survival signaling. Follow up studies using these tools should unravel with more detail the molecular mechanisms mediated by Tns3 not only in immature oligodendrocytes during developmental myelination but also in pathological contexts such as preterm birth dysmyelination, adult demyelination in MS and glioblastoma, this last recently associated with reduced levels of Tns3 (Chen et al., 2017).

## MATERIAL & METHODS

### Animals

All animal procedures were performed according to the guidelines and regulations of the Inserm ethical committees (authorization #A75-13-19) and animal experimentation license A75-17-72 (C.P.). Both males and females were included in the study. Mice were maintained in standard conditions with food and water ad-libitum in the ICM animal facilities. *Tensin3* gene trap mouse line (*Tns3^βgeo^*) was from Su Hao Lo lab (UC Davis, USA). Mice used for ChIP-seq analysis were wild type Swiss obtained from Janvier Labs. *Tns3^flox^* were crossed with *Pdgfra-CreER^T^* (Kang et al., 2010) and *Rosa26^stop-floxed-YFP^*mice to generate *Tns3^flox^*; *Pdgfra-CreER^T^*; *Rosa26^stop-floxed-YFP^* mice line. *Pdgfra-CreER^T^*; *Rosa26^stop-floxed-YFP^* mice were used as controls.

### Generation of *Tns3^Tns3-V5^* knock-in mice

*Tns3^V5^* mice were generated at the Curie Institute mouse facility. Briefly, the Cas9 protein, the crRNA, the tracrRNA and a ssODN targeting vector for the *Tns3* gene had been microinjected into a mouse egg cell, which was transplanted into a C57BL/6J-BALB/cJ female surrogate. Pups presenting HDR insertion of the V5 tag were selected after genotyping.

### Generation of *Tns3^4del^* and *Tns3^14del^* knockout mice

*Tns3^KO^* mice were generated at the ICM mice facility. Briefly, the Cas9 protein, the crRNA, the tracrRNA and a targeting vector for the *Tns3* gene had been microinjected into a mouse egg cell transplanted into a C57BL/6J female surrogate. Pups with NHEJ mutations inducing a gene frameshift were selected after genotyping and Sanger sequencing verification. Finally, only two lines containing indels of 4 and 14 nucleotide deletions were maintained and studied.

### Generation of *Tns3^Flox^* mice and tamoxifen administration

We designed a *Tns3* conditional knockout allele, by flanking with LoxP sites the exon 9 (LoxP-Exon9-LoxP; Fig. S10A). In this *Tns3*-floxed allele (*Tns3^flox^*), Cre-mediated recombination induces a transcription frame shift translated into an early stop codon, leading to a putative small peptide of 109 aa instead of the full length Tns3 protein (1442 aa; Fig. S10B,C). *Tns3^flox^* mice were generated at the Transgenic Core Facility of the University of Copenhagen. The repair template contained homology arms of 771bp and 759bp length and *loxP* sequences flanking Exon 9, and was synthesized by Invitrogen and verified by Sanger sequencing. Two gRNAs were designed at the Transgenic Core Facility that target a DNA sequence in the proximity of each *loxP* site. The gRNAs were designed in a fashion where the insertion of the *loxP* disrupts the targeting site, thus preventing retargeting of the repaired DNA. Mouse embryonic stem cell (mES) method was used for the generation of this mouse model by transfecting ES cells with the repair construct (dsDNA) together with two plasmids – each containing each gRNA. Identification of the positive mES clones was done via a combination of a PCR genotyping and Sanger sequencing confirmation.

Mouse embryonic stem cells (ESCs) were transfected with a plasmid expressing *Cas9, GFP*, and gRNAs flanking *Tns3* exon 9, and a *Tns3*-floxed targeting vector (Fig. S10D), in order to induce CRISPR/Cas9-mediated homologous recombination. After verifying the presence of *Tns3*-floxed allele in *Tns3* locus by Sanger sequencing, positive ESC clones were injected into blastocysts to generate *Tns3*-floxed (*Tns3^flox^*) mice.

Tamoxifen (Sigma, T5648) was dissolved in corn oil (Sigma, C-8267) and injected subcutaneously at 20mg/ml concentration at P7 (30µl) in *Ctrl* and *Tns3-iKO* animals. Brains were then collected at P21.

### Postnatal electroporation

Postnatal brain electroporation (Boutin et al., 2008) was adapted to target the dorsal SVZ. Briefly, postnatal day 2 (P2) pups were cryoanesthetized for 2 min on ice and 1.5 μl of plasmid was injected into their left ventricle using a glass capillary. Plasmids were injected at a concentration of 2-2.5μg/μl. Electrodes (Nepagene CUY650P10) coated with highly conductive gel (Signagel, signa250) were positioned in the dorso-ventral axis with the positive pole dorsal. Five electric pulses of 100V, 50ms pulse ON, 850ms pulse OFF were applied using a Nepagene CUY21-SC electroporator. Pups were immediately warmed up in a heating chamber and brought to their cages at the end of the experiment.

### Demyelinating lesions

Before surgery, adult (2-3months) WT mice were weighed and an analgesic (buprenorphine, 30 mg/g) was administered to prevent postsurgical pain. The mice were anesthetized by anesthetized by induction of isoflurane (ISO-VET). Ocrygel (Tvm) was put on their eye to prevent dryness and lidocaine in cream (Anesderm 5%) was put on the ear bars to prevent pain. After cutting of the skin, a few drop of liquid lidocaine were put to prevent pain. Focal demyelinating lesions were induced by stereotaxic injection of 1µl of lysolecithin solution (LPC, Sigma, 1% in 0.9%NaCl) into the corpus callosum (CC; at coordinates: 1 mm lateral, 1.3 mm rostral to bregma, 1.7 mm deep) using a glass-capillary connected to a 10µl Hamilton syringe. Animals were left to recover in a warm chamber before being returned into their housing cages.

### Tissue processing

Postnatal mouse were transcardially perfused with 15ml (P14) or 25ml (>P21) of 2% PFA freshly prepared from 32% PFA solution (Electron Microscopy Sciences, 50-980-495). Perfused brains were dissected out, dehydrated in 10% sucrose followed by 20% sucrose overnight, and embedded in OCT (BDH) before freezing and sectioning (16µm thickness) in a sagittal plane with a cryostat microtome (Leica).

### Magnetic Assisted Cell Sorting (MACS)

Dissociation of cortex and corpus callosum from mice brain was done using neural tissue dissociation kit (P) (Miltenyi Biotec; ref 130–093-231). Briefly, cortices were dissected from P7, P12, P14 or P21 mice and dissociated using a MACS dissociator (Miltenyi Biotec; ref 130-096-427) followed by filtration through a 70μm cell strainer (Smartstainer; Miltenyi Biotec; ref 130-098-462). Myelin residues were eliminated from P12, P14 and P21 mice cortices during an additional step using the debris removal kit (Miltenyi Biotec; ref 130-090-101). Cells were suspended in a 0.5% NGS solution then incubated with anti-PDGFRα or anti-O4 coupled-beads (Miltenyi Biotec; ref 130-094-543 and 130-096-670). Unbound bead-coupled antibodies were washed away by centrifugation, leaving bound cells which were sorted using MultiMACS Cell24 Separator Plus (Miltenyi Biotec; ref 130-098-637). Sorted cells were either plated in culture plates for *in vitro* cell study or centrifuged at 1200 rpm and used for Western blot analysis or ChIP-seq.

### Immunofluorescence staining and microscopy

Postnatal mouse brain cryosections were dried for 20 minutes at room temperature, before adding the blocking solution (10% normal goat serum (NGS, Eurobio, CAECHVOO-OU) and 0.1% Triton X-100 in PBS) for one hour at room temperature. Primary antibodies were diluted (dilutions indicated in Table 1) in the same blocking solution and incubated on the slices overnight at 4°C. After washing with 0.05% Triton X-100 in PBS, sections were incubated with secondary antibodies conjugated to AlexaFluor-488, AlexaFluor-594 and AlexaFluor-647 (Thermo, 1:1,000). Finally, cell nuclei were labeled with DAPI (1/10000, Sigma-Aldrich®, D9542-10MG), and slices are mounted in Fluoromount-G® (SouthernBiotech, Inc. 15586276).

**Table 1.**

| **Antibodies** |  |  |  |
| --- | --- | --- | --- |
| Rabbit anti Tensin1 | Su Hao Lo, UC Davis | None | 1:100 |
| Rabbit anti Tensin2 | Su Hao Lo, UC Davis | None | 1:100 |
| Rabbit anti Tensin3 | Sassan Hafizi, University of Portsmouth | None | 1:1000 |
| Rabbit anti Tensin3 | Millipore | AB229 | 1:500 |
| Rabbit anti Tensin3 | Thermo-Fisher | PA5-116022 | 1:1000 |
| Mouse monoclonal anti Tensin3 | Santa Cruz Biotech | sc-376367 | 1:500 |
| Rabbit anti Tns3 (C-terminal) | Sigma-Aldrich | SAB4200205 | 1:200 |
| Rabbit anti Tns3 (TN-17) | Sigma-Aldrich | SAB4200416 | 1 :400 |
| Rabbit anti V5 tag | Millipore | AB3792 | 1:2000 |
| Mouse monoclonal anti V5 tag | Invitrogen | R960-25 | 1:1000 |
| Rat anti PDGFRα | BD Biosciences | 558774 | 1:250 |
| mouse monoclonal anti Olig1 | NeuroMab | 75-180 | 1:1000 |
| Mouse monoclonal anti Olig2 | Millipore | MABN50 | 1:500 |
| Mouse monoclonal anti CNPase | Millipore | MAB326R | 1:250 |
| Rabbit anti IP3 receptor 2 (Itpr2) | Millipore | AB3000 | 1:40 |
| mouse polyclonal anti Nkx2.2 | Developmental Studies Hybridoma Bank | None | 1:4 |
| mouse monoclonal anti CC1 (Quaking 7) | Calbiochem | OP80 | 1:100 |
| mouse monoclonal anti MOG | ICM, Paris Hybridoma | AA3 | 1:20 |
| Mouse monoclonal anti Opalin | Santa Cruz Biotech | sc-374490 | 1:500 |
| Chicken polyclonal anti GFP | Aves Labs | GFP-1020 | 1:1000 |
| Rabbit anti-GFP | Life Technologies | A6455 | 1:1000 |
| Mouse anti MCM2 (BM28) | BD Biosciences | 610701 | 1:500 |
| Rabbit anti-p53 | Leica | P53-CM5P-L | 1 :500 |
| Rat anti F4/80 | Abd Serotec | MCA497 | 1:100 |
| Integrin-β1 | Proteintech | 12594-1-AP | 1:100 |
| Mouse monoclonal anti beta-galactosidase | Promega | Z3783 | 1:1000 |
| Chicken anti-GFAP | Aveslabs | 75-240 | 1:1000 |
| HRP-conjugated anti Rabbit | Biorad | 1706515 | 1:5000 |
| HRP-conjugated anti Mouse | Sigma-Aldrich | NA931-1ML | 1:5000 |
| HRP-conjugated anti Rat | Thermo Fisher Scientific | A10549 | 1:5000 |
| rabbit anti-H3K4me3 | Active motif | 39060 |  |
| rabbit anti-H3K27Ac | Active motif | 39034 |  |
| mouse anti-H3K27me3 | Abcam | ab6002 |  |
| rabbit anti-H3K4me1 | Ozyme | 5326T |  |

In Situ Cell Death detection kit (Roche, 12156792910) was used to do Tunel experiment on P21 mouse brains. Briefly, tissues were processed as mentioned above with anti-GFP and anti-CC1 and fixed in fixation solution for 20min at room temperature. After washing, slices were permeabilized for 2 min in permeabilisation solution (0.1% Triton X-100 ; 0.1% sodium citrate) and TUNEL reaction mixture was put on samples for one hour at 37°C. Tissues were then mounted in Fluoromount-G®.

Fixed coverslips were blocked in blocking solution (10% normal goat serum (NGS, Eurobio, CAECHVOO-OU) and 0.1% Triton X-100 in PBS) for 30 minutes at room temperature, incubated in the primary antibodies for 45 minutes at room temperature and washed 3 times in 1x PBS. Secondary antibodies were applied for 45 minutes at room temperature and washed 3 times in 1X PBS. Coverslips were then incubated with DAPI solution for 5 minutes at RT. A final washing was done before mounting the coverslips on slides to be visualized under the microscope.

Immunofluorescence was visualized with Zeiss® Axio Imager.M2 microscope with Zeiss® Apotome system. Pictures were taken as stacks of 5–10µm with 0.5µm between sections. Image acquisition and processing are achieved by ZEN Microscopy and imaging Software, Z-projections and orthogonal projections were done in ImageJ and processed with Adobe Photoshop. Figures were made using Adobe Illustrator.

### Western blot

Proteins from MACsorted cells were extracted during 30 minutes at 4°C in RIPA buffer (ThermoFisher; 50 µL per million cells, 89901) supplemented with Halt™ Protease Inhibitor Cocktail (100X; ThermoFisher, 87786). Protein concentration in the supernatant was estimated using the Pierce Detergent Compatible Bradford Assay Kit (Thermofisher, 23246). For each Western Blot, we used 50µg of proteins denaturated for 10 minutes at 95°C with added β-mercaptoethanol (from 24X stock) and BoltTM LDS Sample Buffer (4X) (ThermoFisher, B0007). Sodium dodecyl sulfate-polyacrylamide gel electrophoresis (SDS-PAGE) was performed using precast 4-12% polyacrylamide gradient gels (ThermoFisher, NW04122BOX), submerged at 4°C in Bolt™ MOPS SDS Running Buffer (ThermoFisher, B0001) using Mini Gel Tank and Blot Module Set (ThermoFisher, NW2000). Precision Plus Protein™ All Blue protein standards (BioRad, 1610373EDU) were run alongside the samples as a protein migration control. Proteins were separated for 90 minutes at 90V, after which gels were transferred onto Amersham Protran 0.2 µm nitrocellulose membrane (Dutscher, 10600001) immersed at 4°C in NuPAGE Transfer Buffer (ThermoFisher, NP0006-1) for 90 minutes at 60V. Following transfer, membranes were incubated for 1h in TBS-T, 10% dry milk to aid blocking of non-specific binding by the antibodies. Primary antibodies diluted in TBS-T were incubated with the membrane overnight at 4°C with shaking. After three washes in TBS-T, membranes were incubated with HRP-conjugated secondary antibodies diluted in TBS-T for 1h at 4°C with shaking, then developed using Pierce™ ECL Western Blotting Substrate (ThermoFisher, 32109) and imaged with the ChemiDoc™ Touch Imaging System (BioRad, 1708370). Western blot detection of actin was performed as loading control.

### Videomicroscopy

Tamoxifen was administered to P5 Tns3^flox^; Pdgfrα-CreER^T^; Rosa26^stop-floxed-YFP^ and Tns3^flox^; Pdgfrα-WT; Rosa26^stop-floxed-YFP^ littermates. Brains were dissected out at P7 in order to MACSort OPCs using an anti-PDGFRα antibody coupled to magnetic beads. OPCs were plated in poly-L-ornithine (sigma, P4957) coated µ-Slide 8 Well Glass Bottom slide (Ibidi, 80827) at 40,000 cells/mm2 in OPC proliferative medium : DMEM/F12 (life technology, 31331028), 5mM HEPES buffer (life technology, 15630056), 0.6% glucose (Sigma, G8769), 1x penicillin/streptomycin (life technology, 15140122), N2 supplement (life technology, 17502048), B27 supplement (life technology, 17504044), 20ng/µl EGF (Peprotech, AF-100-15), 10ng/µl FGF-basic (Peprotech, 100-18B), 10ng/µl PDGF-AA (PeproTech, 100-13A) and 20μg/ml Insulin (Sigma, I6634). After 3 days of proliferation, medium was replaced by growth factor depleted medium. Cell differentiation was tracked for 3 days using time-lapse video recording. Cells were put in to a videomicroscope (Zeiss AxioObserver 7, provided by ICM-quant and CELIS facilities) with a humidified incubator at 37°C with a constant 5% CO2 supply. Images for both FITC and bright field were acquired every 10 minutes.

### Chromatin immunoprecipitation (ChIP)

ChIP-seq assays were performed as described previously (Marie et al; 2018), using iDeal ChIP-seq kit for Transcription Factors (Diagenode, C01010055). Briefly, O4^+^ MACSorted cells were fixed in 1% formaldehyde (EMS, 15714) for 10 min at room temperature and the reaction was quenched with 125 mM glycine for 5 min at room temperature. Lysates were sonicated with a Bioruptor Pico sonicator (Diagenode, total time 8 min) and 4μg of antibodies were added to sheared chromatin (from 4 million cells for Olig2 and from 1 million cells for histone marks) and incubated at 4°C overnight on 10 rpm rotation. Antibodies used were: mouse anti-Olig2 antibody (Millipore, MABN50), rabbit anti-H3K4me3 antibody (Active motif, 39060), rabbit anti-H3K27Ac antibody (Active motif, 39034), rabbit anti-H3K4me1 antibody (Ozyme, 5326T), mouse anti-H3K27me3 antibody (Abcam, ab6002). Mock (Rabbit IgG) was used as negative control. Chromatin-protein complexes were immunoprecipitated with protein A/G magnetic beads and washed sequentially according to the manufacturer (Diagenode, C01010055). DNA fragments were then purified using IPure beads v2 (Diagenode, C01010055). Input (non-immunoprecipitated chromatin) was used as control in each individual experiment.

The ChIP-seq libraries were prepared using ILLUMINA Truseq ChIP preparation kit and sequenced with ILLUMINA Nextseq 500 platform.

### ChIP-seq analysis

All ChIP-seq analysis were done using the Galaxy Project (https://usegalaxy.org/). Reads were trimmed using Cutadapt (--max-n 4) and Trimmomatic (TRAILING 1; SLIDINGWINDOW 4 and cutoff 20; LEADING 20; MINLEN 50), and mapped using Bowtie2 onto mm10 mouse reference genome (-X 600; -k 2; --sensitive). PCR-derived duplicates were removed using PICARD MarkDuplicates. Bigwig files were generated with bamCoverage (binsize=1). Peak calling was performed using MACS2 callpeak with Input as control and with options: --qvalue 0.05; -- nomodel; --keep-dup 1; --broad (only for histone marks). Blacklisted regions were then removed using bedtools Intersect intervals.

Visualization of coverage and peaks was done using IGV ((Robinson et al., 2011); http://software.broadinstitute.org/software/igv/home). Intersection and analysis of bound genes were done using Genomatix (https://www.genomatix.de/). Chd7, Chd8 and Mock ChIP-seq datasets are from (Marie et al., 2018). Heatmap was done using R (4.0) using pheatmap package. GO analysis was done using Enrichr GO Biological Process 2021.

Two replicates were done for Olig2, with one of them of better quality (53,960 peaks for replicate 1 and 14,242 peaks for replicate 2). Only the peaks found in both replicates (6,781) and the peaks from replicate 1 found in regulatory elements (13948) were considered (16578 in total). Three replicates were done for H3K4me3, two replicates were done for H3K27me3 and one replicate was done for H3K27Ac and H3K4me1. Intersection of these datasets was done using bedtools Intersect intervals.

Peaks overlapping with regions between 1000bp upstream of transcription start site (TSS) and 10bp downstream of TSS were identified as “promoters” (Genomatix). “Active promoters” were represented by peaks for H3K4me3 and H3K27Ac. “Repressed promoters” were represented by peaks for H3K27me3 and no active marks. “Poised promoters” were represented by peaks for H3K4me1 and no active or repressed mark. Regions outside promoters containing histone marks were considered as “enhancers”. “Active enhancers” were represented by peaks for H3K27Ac. “Repressed enhancers” were represented by peaks for H3K27me3 and no active marks. “Poised enhancers” were represented by peaks for H3K4me1 and no active or repressed mark. Genes were considered associated if the peaks were present in the promoter or within a range of 100kb from the middle of the promoter and the gene expression was medium to high (“active”), low (“poised”) or not (“repressed”) expressed (based on control RNA-seq dataset in GSE116601).

### RNAseq analysis

Raw data were downloaded from GEO datasets GSE107919 and GSE116601 and processed throught the Galaxy Project (https://usegalaxy.org/) using RNAstar for alignment on mm10 reference genome and featureCounts to obtain counts. CPM (count per million), FPKM and statistical analysis were done with R (4.0) using edgeR quasi-likelihood pipeline.

Using control RNA-seq dataset in GSE116601, genes were classified based on their expression as not (below first quartile), low (between first quartile and mean), medium (between mean and third quartile) and high (above third quartile).

### scRNAseq analysis

Counts per gene were downloaded from GEO datasets GSE75330 and GSE95194, and processed in R (4.0) using the following packages: *Seurat* (3.0) for data processing, *sctransform* for normalization, and *ggplot2* for graphical plots. Seurat objects were first generated for each dataset independently using *CreateSeuratObject* function (min.cells = 5, min.features = 100). Cell neighbors and clusters were found using *FindNeighbors* (dims = 1:30) and *FindClusters* (resolution = 0.4) functions. Clusters were manually annotated based on the top 50 markers obtained by the *FindAllMarkers*, adopting mainly the nomenclature from Marques 2016. Using the *subset* function, we selected only the clusters containing neural progenitors and oligodendroglia cells. Using the *merge* function, we combined both oligodendroglial datasets into a single Seurat object (OLgliaDevP) containing 5516 cells. The new object was subjected to *NormalizeData*, *FindVariableFeatures*, *ScaleData*, *RunPCA*, and *RunUMAP* functions with default parameters. Different OPC clusters were fused into a single one keeping apart the cycling OPC cluster. For DimPlots and Dotplots, clusters were ordered by stages of oligodendrogenesis from neural stem cells (NSCs) to myelinating OLs. R script is provided as supplementary file.

### *Tns3^βgeo^* mice analysis

To explore the role of Tns3 in OL differentiation, we first analyzed a *Tns3* gene trap mouse line (*Tns3^βgeo^*) previously studied outside the CNS (Chiang et al., 2005), where the *βgeo* cassette is inserted after *Tns3* exon 4 (Fig. S7A) driving *LacZ* transcription and by inserting a stop poly-A sequence, predicted to be a *Tns3* loss-of-function mutation. Despite the original report of postnatal growth retardation in *Tns3^βgeo/βgeo^* mice, these mice were kept in homozygosity for several generations in C57BL/6 genetic background (Su-Hao Lo, UC Davis). We thus analyzed the impact in oligodendrogenesis in the postnatal brain of *Tns3^βgeo^* animals. We first immunodetected βgalactosidase in OLs (Olig2^+^/CC1^+^ and Olig2^+^/PDGFRα^−^ cells) of *Tns3^βgeo^* postnatal brains at P21 (Fig. S7B,C), paralleling our characterization of Tns3 expression with V5 and Tns3 antibodies. We then quantified the density of PDGFRα^+^ OPCs or CC1^+^ oligodendrocytes in *Tns3^βgeo/βgeo^* and *Tns3^βgeo/+^* littermates at P21, finding similar number of OPCs and oligodendrocytes in two main white matter areas (corpus callosum and fimbria; Fig. S7D,D’,E). Moreover, quantification of three different stages of oligodendrocyte differentiation (iOL1, iOL2, and mOL) by Olig2/CC1/Olig1 immunofluorescence did not reveal changes in the rate of oligodendrocyte differentiation (proportion of each stage) in *Tns3^βgeo/βgeo^* mice compared to control littermates (Fig. S7F,F’,G). We verified the homozygosity of *Tns3^βgeo^* allele in *Tns3^βgeo/βgeo^* mice by PCR amplification from genomic DNA of P21 brains finding that primers recognizing Intron 4 and βgeo only produced PCR amplicons in *Tns3^βgeo/βgeo^* mice but not when using Intron 4 flanking primers that only produced PCR amplicons in wild type mice; Fig. S7H). We then checked for *Tns3* full-length transcripts using cDNA generated from P21 brains, and to our surprise, primers flanking Exons 17 and 31 were similarly amplified from cDNA of *Tns3^βgeo/βgeo^* and wild type brains (Fig. S7I), suggesting that in the brain of *Tns3^βgeo/βgeo^* mice *Tns3* full-length transcripts coding for Tns3 protein are still produced. Altogether, these results suggested that, unlike in other tissues (Chiang et al., 2005), the *Tns3^βgeo^* allele does not lead to *Tns3* loss-of-function in the brain, likely through the generation of alternative spliced *Tns3* variants, and is thus not suitable for assessing *Tns3* function in the CNS.

### Analyses of *TNS3* alleles in the human population based in gnomAD project

To assess whether TNS3 is potentially required during human development, we explore for the presence of *TNS3* gene variants in the human population using the gnomAD database containing 125,748 exomes and 15,708 whole-genome sequences from unrelated individuals (Karczewski et al., 2020; Lek et al., 2016). Homozygous predicted loss-of-function (pLoF) alleles of *TNS3* were not found, and heterozygous pLoF were greatly below the expected frequency (0.1 observed/expected ratio, with 90% confidence interval of 0.05–0.19; and LOEUF of 0.19; Fig. S8A; https://gnomad.broadinstitute.org), meaning that heterozygous loss-of-function variants of TNS3 causes ∼80% developmental mortality, a rate similarly high to key neurodevelopmental genes such as *SOX10* (LOEUF=0.21; Fig. S8B), *CHD7* (LOEUF=0.08; Fig. S8C), and *CHD8* (LOEUF=0.08; Fig. S8D), contrary to less broadly required factors such as *NKX2-2* (LOEUF=0.67; Fig. S8E) and *OLIG1* (LOEUF=1.08; Fig. S8F). Therefore, *TNS3* loss-of-function variants are badly tolerated in both mouse and human development.

### CRISPR/Cas9 tools development

CRISPOR software (http://crispor.tefor.net/) was used to design gRNAs with predicted cutting efficiency and minimal off-target and PCR amplification primers. The validation of *Tns3*-targeting CRISPR/Cas9 system was performed in 3T3 cells by transfection with Lipofectamine 3000 of *PX459* plasmids containing 4 different sgRNA sequences. After 2 days incubation, puromycin was added to medium for 4 days allowing survival of cells containing the *PX459* plasmid. Three days after proliferation in fresh medium without puromycin, DNA was extracted using DNeasy blood & tissue kit (Qiagen). The target DNA for 5’ *Tns3* region was amplified by PCR using primers with the following sequences: Forward: 5’-AGG TGG CCT TCA GCT CAGT-3’, Reverse: 5’-GCT ATC ATC CCC ACT CAC CA-3’; annealing temperature of 64°C, with the PCR product expected to be 326bp. DNA from 3’ *Tns3* target region was amplified using primer with the following sequences: Forward: 5’-CCA GTC AGT GGT GAC ATT GTTT-3’, Reverse: 5’-ACT GTT CCC AGG TTG CTA TCAT-3’), annealing temperature of 58°C, with the PCR product expected to be 419bp.

Cutting efficiency of sgRNA was verified by T7 endonuclease I, following the beta protocol of IDTE synthetic biology for amplification of genomic DNA and detecting mutations (Fig. S9C), and using PAGE (Fig. S9D). In order to generate plasmids that will insert CRISPR tools into the genome of the transfected cells and lead to permanent expression of the targeting tools, the *PX458* (GFP) or *PX459* (Puromycin) plasmids were subcloned into a *Tol2*-containing sequence backbone (obtained from *Tol2-mCherry* expressing plasmid kindly provided by Jean Livet, Institut de la Vision, Paris).

### *Tns3^4del^* and *Tns3^14del^* mice generation by CRISPR and analyses

To generate new *Tns3* knockout mouse using CRISPR/Cas9 technology, by introducing loss-of-function mutations (indels) at the beginning of *Tns3* full-length coding sequence. We generated CRISPR integrative plasmids (Fig. S9A) driving Cas9 expression and gRNAs targeting *Tns3* Exon-6 at the levels of the first coding ATG, using as control, plasmids without the *Tns3*-targeting sequence of the gRNA. Strong cutting efficiency of two gRNAs was validated by lipofection of neural progenitors (Fig. S9B-F). We then used these optimized tools to induce CRISPR-mediated *Tns3* mutations in mouse zygotes, generating and characterizing two mouse lines having small deletions (4-deletion and 14-deletion) after the first coding ATG of *Tns3* (Fig. S7J), expected to cause frame shifts leading to *Tns3* loss-of-function. Remarkably, homozygous animals were found in reduced numbers compared to mendelian ratios with many of them dying during embryonic development (Fig. S7K) with most homozygous animals showing major growth retardation by the second postnatal week compared with their littermates (Fig. S7L), similar to the original report of *Tns3^βgeo^* mice (Chiang et al., 2005). Furthermore, we could still immunodetect Tns3 protein in CC1^+^ oligodendrocytes of these homozygous mice at P21 with at least two different Tns3 antibodies (Fig. S7M,N), and detect *Tns3* exons corresponding to *Tns3* full-length transcript by qPCR (Fig. S7O). Further analysis of these mice was prevented by the Covid-19 lockdown leading to the loss of these *Tns3* knockout mouse lines. Altogether, these results suggest that mice carrying constitutive *Tns3* loss-of-function mutations seems to escape the full *Tns3* loss-of-function in the brain, by generating alternative spliced variants containing the main *Tns3* full-length exons, and thus we considered these animals not suitable to study *Tns3* function in oligodendrogenesis.

### RNA sequencing and analysis

Cortices from 3-4 animals for each group were dissected and frozen in liquid nitrogen for further processing. Total RNA was isolated with the TRIzol Reagent protocol (TermoFisher) from spinal cords and RNeasy Mini Kit (Qiagen) according to instructions of the provider. The RNA-seq libraries were prepared using either the NEBNext Ultra II Directional RNA Library Prep Kit (NEB) and sequenced with the Novaseq 6000 platform (ILLUMINA, 32*106 100bp pair-end reads per sample). Quality of raw data was evaluated with FastQC. Poor quality sequences were trimmed or removed with fastp tool, with default parameters, to retain only good quality paired reads. Illumina DRAGEN bio-IT Plateform (v3.6.3) was used for mapping on mm10 reference genome and quantification with gencode vM25 annotation gtf file. Library orientation, library composition and coverage along transcripts were checked with Picard tools. Following analysis were conducted with R software. Data were normalized with edgeR (v3.28.0) bioconductor packages, prior to differential analysis with glm framework likelihood ratio test from edgeR package workflow. Multiple hypothesis adjusted p-values were calculated with the Benjamini-Hochberg procedure to control FDR. For the differential expression analyses, low expressed genes were filtered, sex was used as covariable (when relevant) and the cut-offs applied were FDR < 0.05. Finally, gene ontology (GO) enrichment analysis of biological processes of the differentially expressed genes (DEGs) was conducted with clusterProfiler R package (v3.14.3).

### Data Resources

Raw data files have been deposited in the NCBI Gene Expression Omnibus under accession number GEO GSE203295.

#### Contact for Reagent and Resource Sharing

Further information and requests for reagents may be directed to, and will be fulfilled by the corresponding author Carlos Parras.

## ACKNOWLEDGMENTS

We thank Dwight Bergles for the *PDGFRα::CreER^T^* mice and Jean Livet for Tol2-mCherry plasmid. Mathilde Bertrand for sharing ChIP-seq analysis pipeline. All animal work was conducted at the ICM PHENOPARC Core Facility. Data generated relied on ICM Core Facilities: PHENO ICMice, iGenSeq, DAC, iVector, CELIS, Histomics, and ICM Quant, and we thank all personnel involved for their contribution and help. The Core Facilities were supported by the “Investissements d’avenir” (ANR-10-IAIHU-06 and ANR-11-INBS-0011-NeurATRIS) and the “Fondation pour la Recherche Médicale”. This work was supported by funding by grants from the National Multiple Sclerosis Society (NMSS RG-1501-02851), and the Fondation pour l’Aide à la Recherche sur la Sclérose en Plaques (ARSEP 2014, 2015, 2018, 2019, 2020). E.M, H.H, and C.M. were supported by funding from Sorbonne Université. C.M. was also supported by Fondation pour la Recherche Médicale (FRM, FDT20160435662) and ARSEP grant 2018-2020.

---

**Figure S1.**
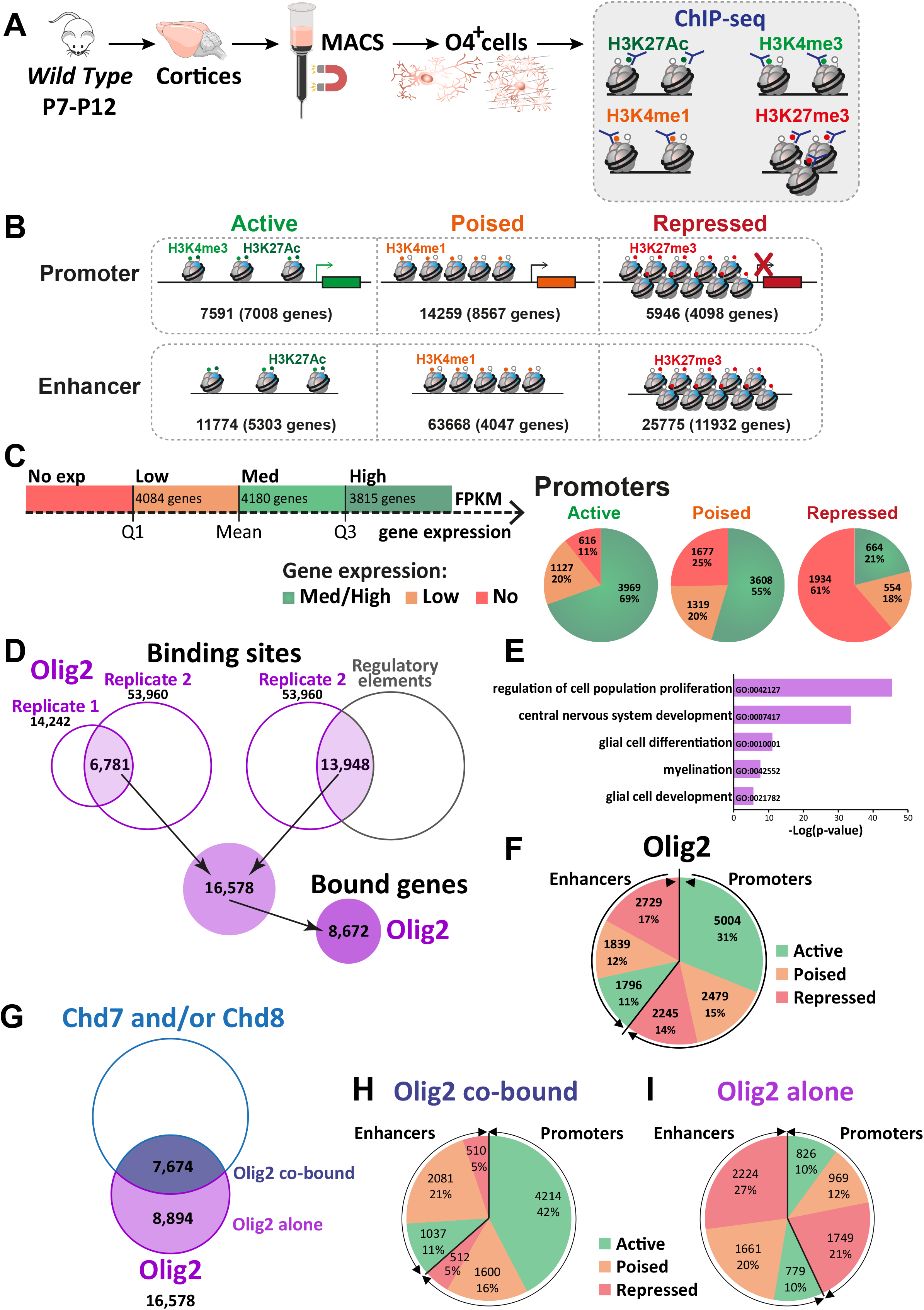
*Tns3* is a target gene of Olig2 and Chd7/8 regulators of oligodendrocyte differentiation. **(A)** Scheme representing MACSorting of O4^+^ cells from wild type cortices followed by ChIP-seq for histone marks H3K4me3, H3K27Ac, H3K4me1, and H3K27me3. **(B)** Schematic representing gene regulatory elements in O4^+^ cells as active, poised, and repressed, according to their histone marks. **(C)** Left, diagram representing the classification of gene expression (data from Marie et al, 2018). Right, Pie chart representing percentage of medium-to-high, low or not expressed genes having an active, poised, and repressed promoter in O4^+^ cells. Note the high percentage of medium-to-high expressed genes having active marks in their promoter and the high percentage of not expressed genes having repression marks in their promoter. **(D)** Venn diagram representing the method used to determine Olig2 binding peaks. As the quality of the peaks in replicate 2 was better than in replicate 1, we considered peaks common in both replicate and the peaks of replicate 2 that were present in regulatory elements (regions with histone marks). 16,578 peaks were found corresponding to 8,672 genes bound by Olig2. **(E)** Gene ontology category found enriched in Olig2 bound genes. **(F)** Pie chart representing the number and repartition of Olig2 binding sites in regulatory regions with different activity histone marks. Note that Olig2 binds in both promoters (60%) and enhancers (40%), and that it is found in either active, poised or repressed regions. **(G)** Venn diagram representing the amount of Olig2 binding sites which are also bound by Chd7 and/or Chd8 (7,674; Olig2 co-bound) or not (8,894; Olig2 alone). **(H-I)** Pie chart representing the number and repartition of Olig2 co-bound **(H)** or alone **(I)** sites in regulatory regions with different activity histone marks. Note that 42% Olig2 co-bound sites are in active promoters, whereas 48% of Olig2 alone sites are in repressed promoter or enhancers.

**Figure S2.**
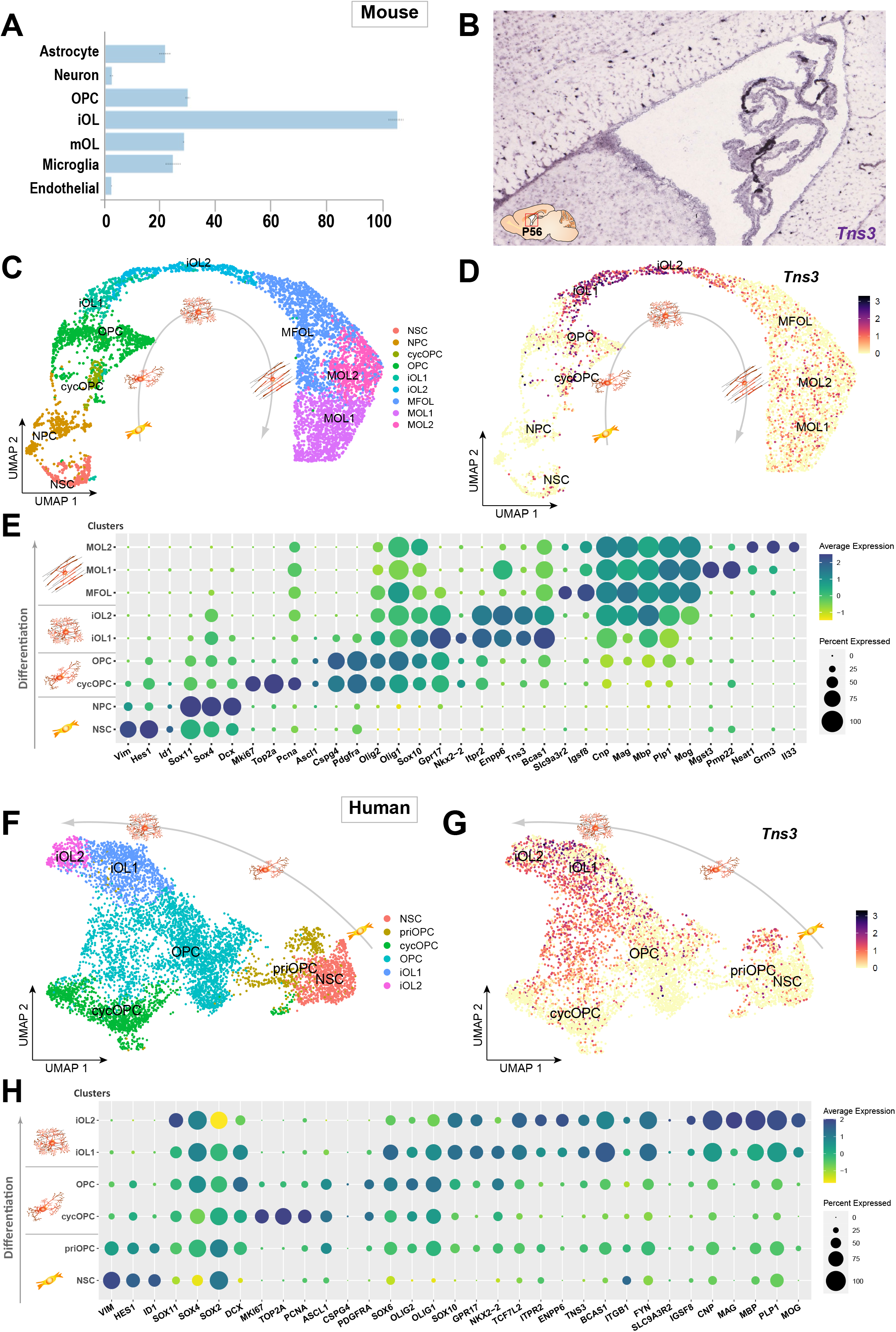
Strong expression of *Tns3* transcripts in mouse & human immature oligodendrocytes. **(A)** Barplot of Tns3 mRNA transcript levels (FKPM) in postnatal brain cell types (Zhang et al., 2014; brainrnaseq.org). **(B)** In situ hybridization in sagittal section of the adult (P56) mouse brain at the level of the lateral ventricles showing *Tns3* transcript expression in sparse white matter cells (Allen brain atlas, portal.brain-map.org). **(C)** UMAP representation of neural progenitors and oligodendroglial cells extracted and integrated using Seurat from scRNA-seq datasets (Marques et al., 2016; 2018), representing different clusters corresponding to different oligodendroglial stages of NSCs to mOLs. **(D)** Feature plot representing relative expression levels of *Tns3* transcript. **(E)** Dot plot representing the transcript expression of key markers for each cell stage/subtype and key oligodendroglial factors in the different clusters showing the predominant expression of *Tns3* in iOL1s and iOL2s clusters, similar to *Enpp6* and *Itpr2*. NSC, neural stem cells; NPC, neural progenitor cells; cycOPC, cycling OPC; MFOL, myelin forming OL; MOL1/2, myelinating OL 1/2. **(F-H)** *TNS3* expression in human oligodendroglial cells differentiated from iPSCs by scRNA-seq (Chamling et al., 2021), showing *TNS3* high levels of expression in immature oligodendrocytes. **(F)** UMAP representation showing six clusters (NSC: neural stem cells, cycOPC: cycling OPCs, OPCs, and two iOL stages, iOL1 and iOL2, similar to mouse oligodendroglia). **(G)** Feature plot showing *TNS3* expression levels. **(H)** Dotplots showing *TNS3* expression pattern together with key markers of each cluster and regulators of oligodendrogenesis.

**Figure S3.**
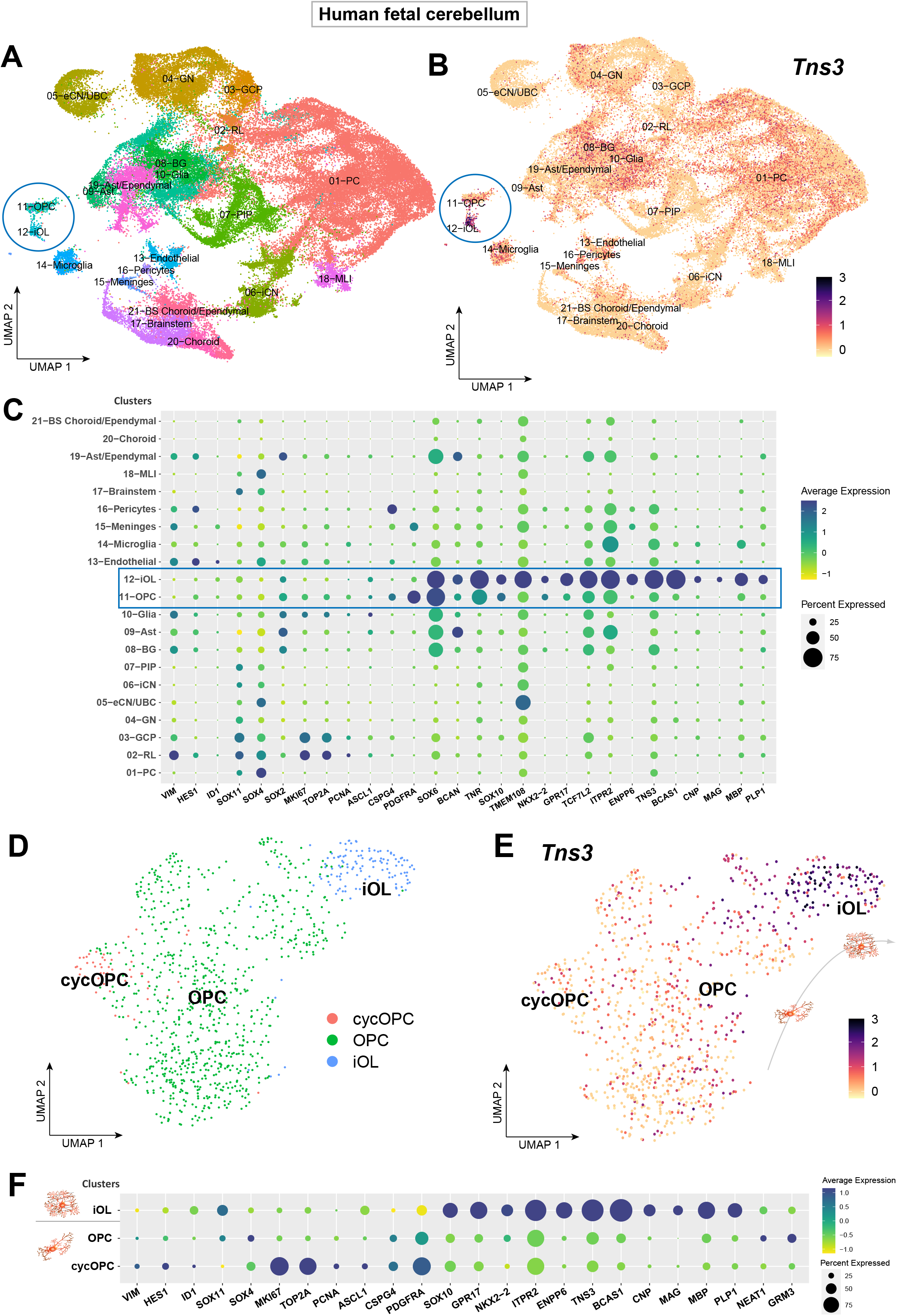
Strong expression of *Tns3* transcripts in immature oligodendrocytes of midterm human cerebellum. **(A)** UMAP representation of different cell-type cluster found in human fetal cerebellum (GW 9-22), including OPCs and iOLs clusters (blue circle). **(B)** Feature plot representing relative expression levels of *Tns3* transcript, presenting high levels in iOL cluster cells **(C)** Dotplots representing the transcript expression of key markers for each cell stage/subtype and key oligodendroglial factors in the different clusters showing the predominant expression of *TNS3* in iOLs clusters (OPC and iOL clusters highlighted by blue rectangle). **(D)** UMAP representation of new clustering of oligodendroglial cells (11-OPC and 12-iOL clusters) showing OPCs, cycling (cyc) OPCs and iOLs. **(E)** Feature plot representing relative expression levels of *TNS3* transcript showing highest levels in iOL cells. **(F)** Dotplots showing *TNS3* expression pattern together with key markers of each cluster and regulators of oligodendrogenesis, showing similar expression pattern of *TNS3* and other suggested iOL markers (including *ITPR2, ENPP6*, and *BCAS1*).

**Figure S4.**
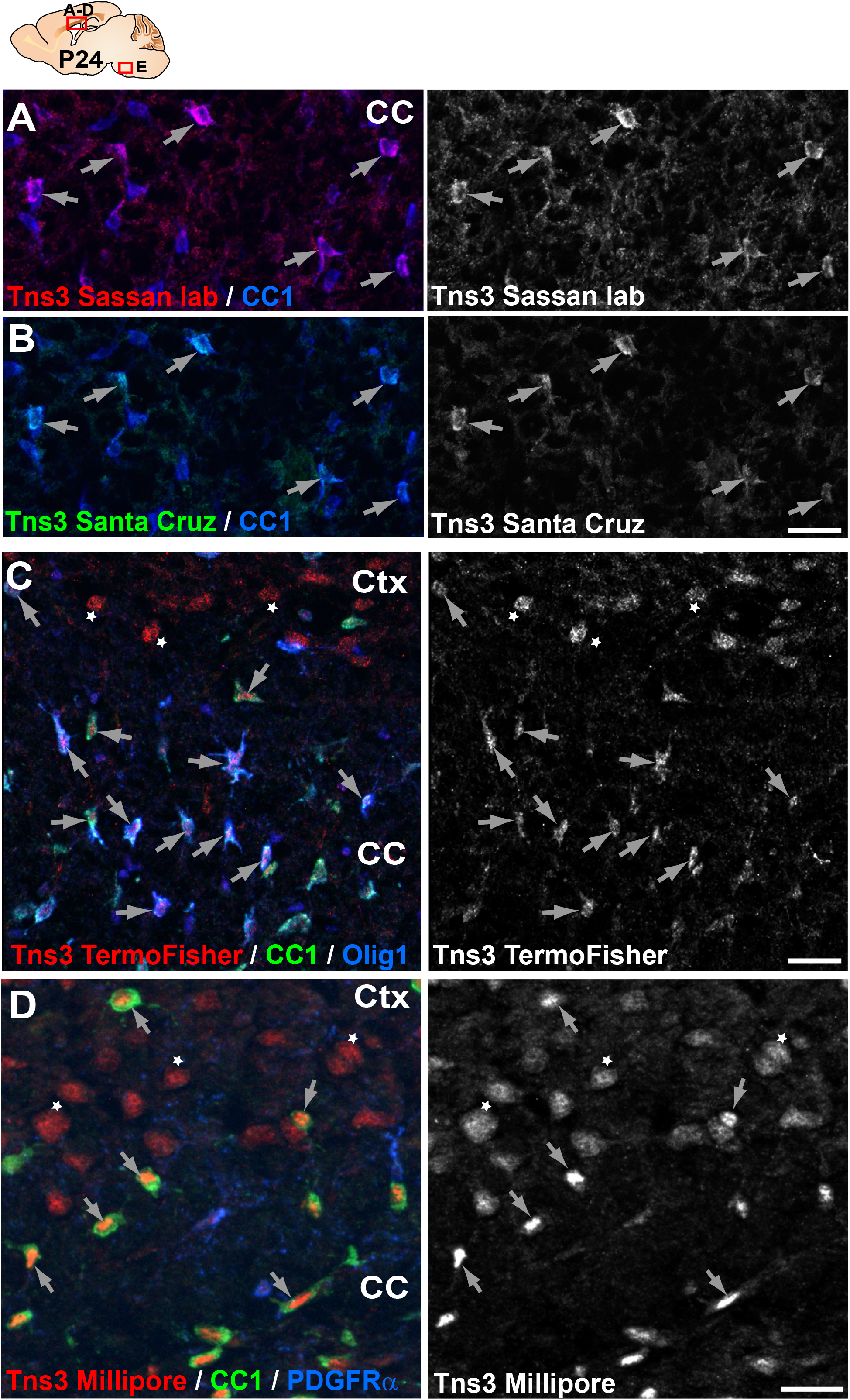
Immunodetection of Tns3 protein in immature oligodendrocytes of the postnatal brain. **(A-D)** Tns3 expression in CC1^+^ OLs (arrows) at the level of the corpus callosum (CC) of P24 mice, using a homemade antibody from Sassan Hafizi lab (University of Portsmouth) (A), and commercial antibodies from Santa Cruz (B), ThermoFisher (C), and Millipore (D). Note that Millipore antibody is the only one to show clear nuclear localization in OLs. ThermoFisher and Millipore antibodies also recognize a nuclear signal in cortical neurons (C-D, stars), not found with other Tns3 antibodies. Scale bar, 20 μm.

**Figure S5.**
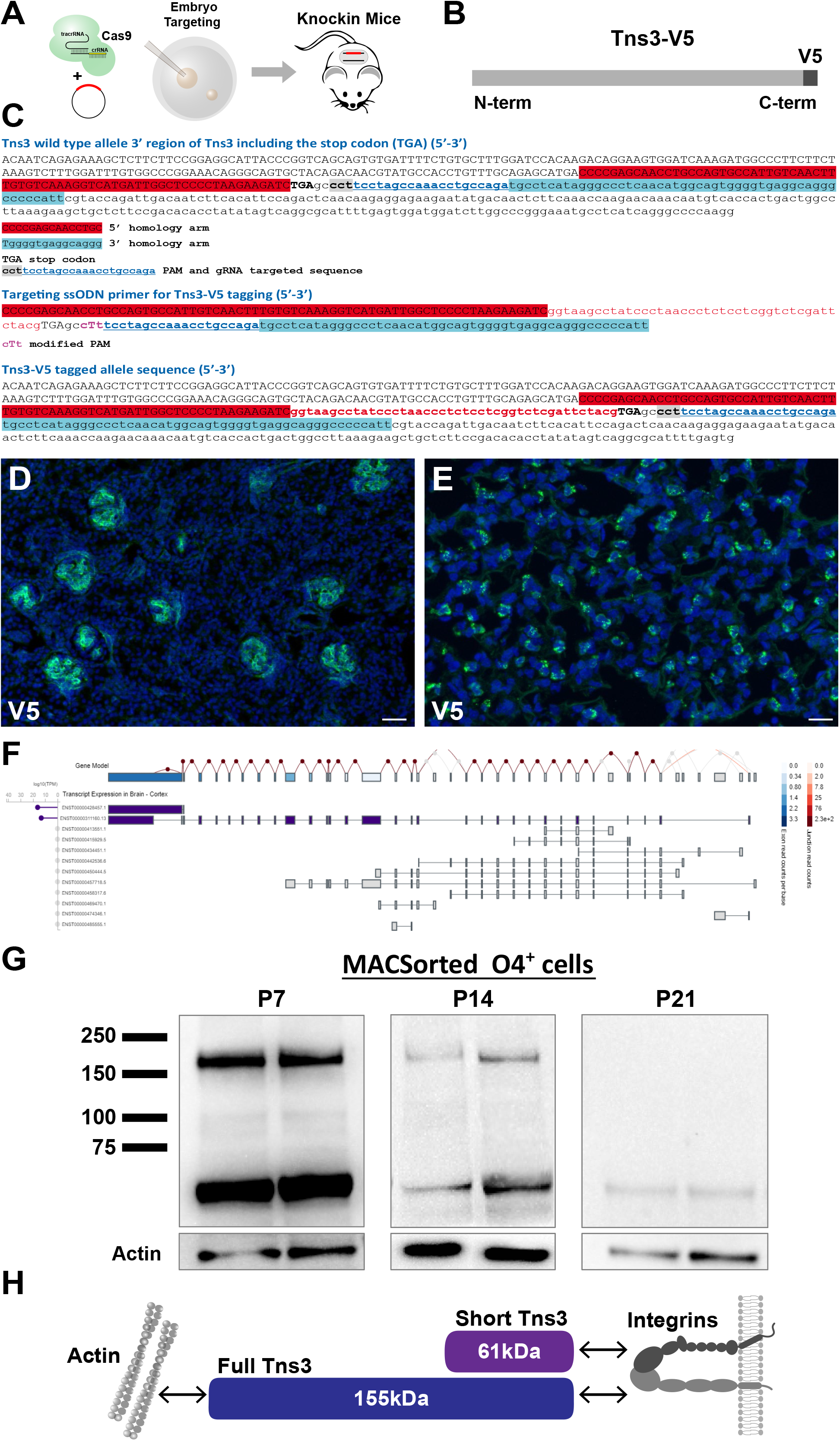
Generation of *Tns3^Tns3-V5^*knock-in mice. **(A)** Scheme of V5 tag knock-in strategy in mice. **(B)** Schematic of Tns3 protein with a V5 tag at the C-terminal. **(C)** Nucleotide sequence of a wild type mice *Tns3* 3’ region (Upper), of the targeting ssODN vector (Middle) and of the 3’ region of the first Tns3^Tns3-V5^ knock-in mice generated (Lower). **(D,E)** Immunostaining using V5 antibody in kidney (D) and lung (E) of P14 *Tns3^Tns3-V5^* mice. Note the expression of Tns3 in kidney glomeruli and lung alveolar cells **(F)** Schematic of the two main *TNS3* isoforms exon expressed in the human brain (gtexportal.org/home/gene/TNS3). A dark color is associated with higher expression. **(G)** Western blot of V5 antibody in O4^+^ MACSorted cells from P7 OPCs/iOLs (left), P14 iOLs (middle) and P21 OLs (right) of *Tns3^Tns3-V5^* mouse brains. Bottom, Actin loading control. Note the presence of the full-length isoform only in OPC/iOLs. **(H)** Schematic of Tns3 protein brain isoforms and their interactions with actin and integrins.

**Figure S6.**
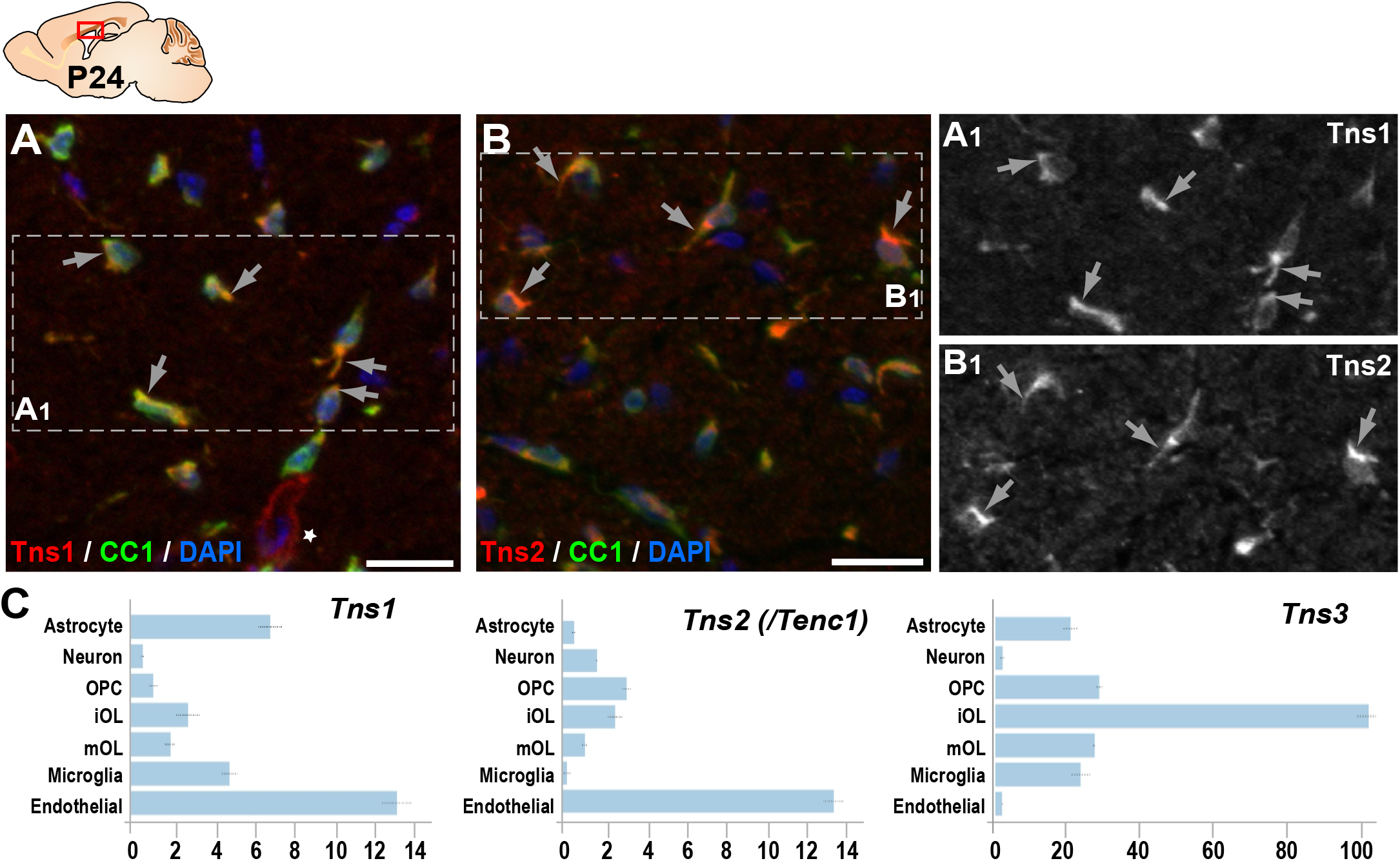
Tns1 and Tns2 proteins are detected at low levels in immature oligodendrocytes. Immunofluorescence in sagittal section of P24 mouse brain. Tns1 **(A)** and Tns2 **(B)** expression in CC1^+^ OLs of the corpus callosum. Tns1 expression is cytoplasmic and excluded from OL nucleus **(A_1_)**, as is Tns2 **(B_1_)**. **(C)** Tns1, Tns2 and Tns3 mRNA expression in postnatal brain (brainrnaseq.org). Note that Tns3 is the main Tensin expressed in oligodendroglia. Scale bar, 20 μm.

**Figure S7.**
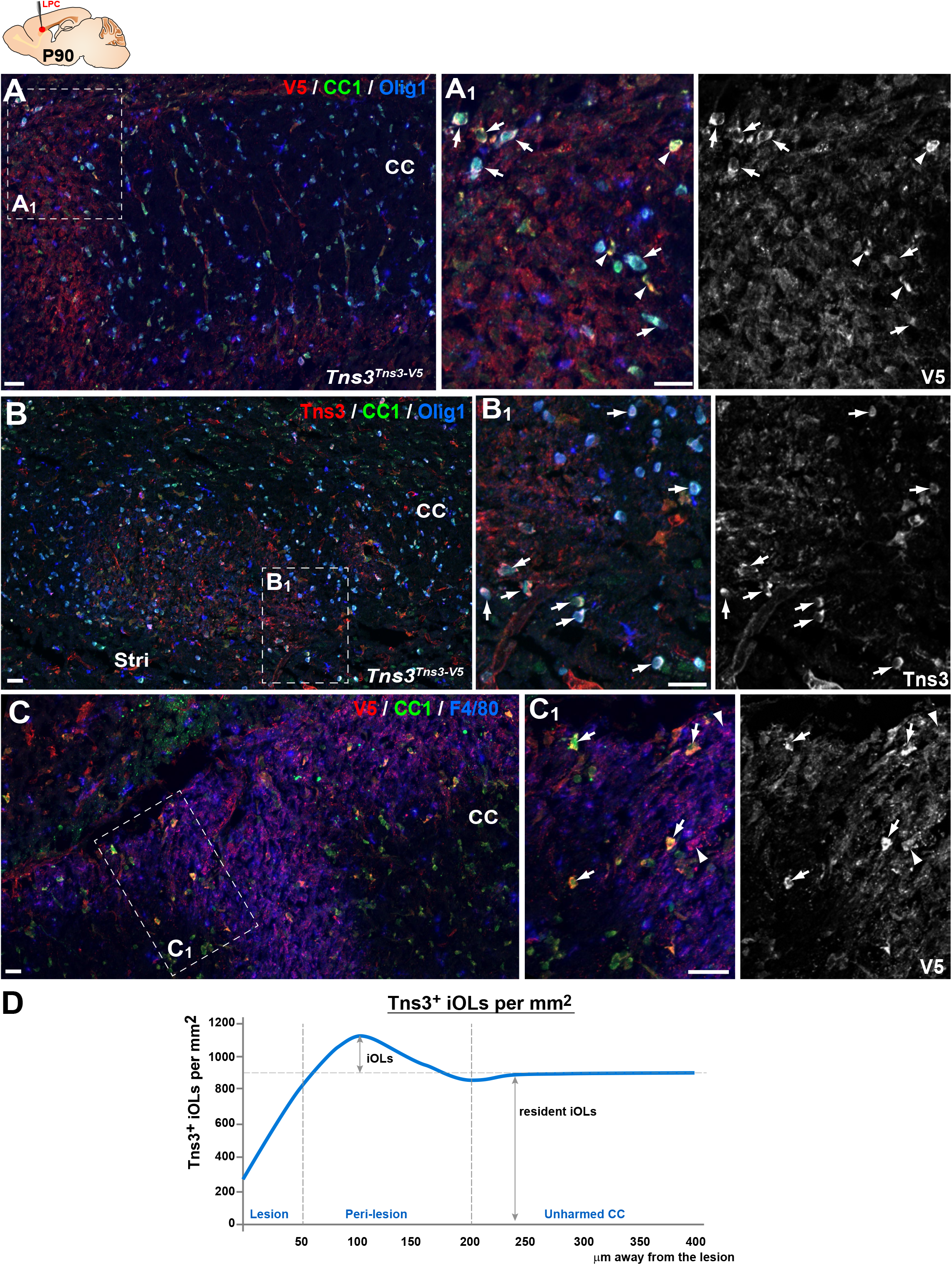
Tns3 is expressed in newly formed oligodendrocytes during adult brain remyelination. **(A,B)** Tns3-expressing cells in the CC of adult (P90) mice, 7 days after chemically induced demyelination, showing strong Tns3 expression in CC1^+^/Olig1^-^ iOL1s (arrowheads) and CC1^+^/Olig1^cytoplasmic^ iOL2s (arrows) around the lesion using either anti-V5 antibody in *Tns3^Tns3^*^-V5^ mice (A-A_1_) or anti-Tns3 antibody in wild type mice (B-B_1_). Note the absence of CC1^+^/Olig1^+^ OLs in the lesion. **(C-C_1_)** Tns3 expression in CC1^high^ iOLs (arrows) around the lesion and in some F4/80^+^ microglia (arrowheads) in the lesion area. A_1_, B_1_ and C_1_ are higher magnification of corresponding insets (squares). **(D)** Quantification of the Tns3^+^ OLs density each 50μm distance away from the lesion. Scale bar, 20 μm.

**Figure S8.**
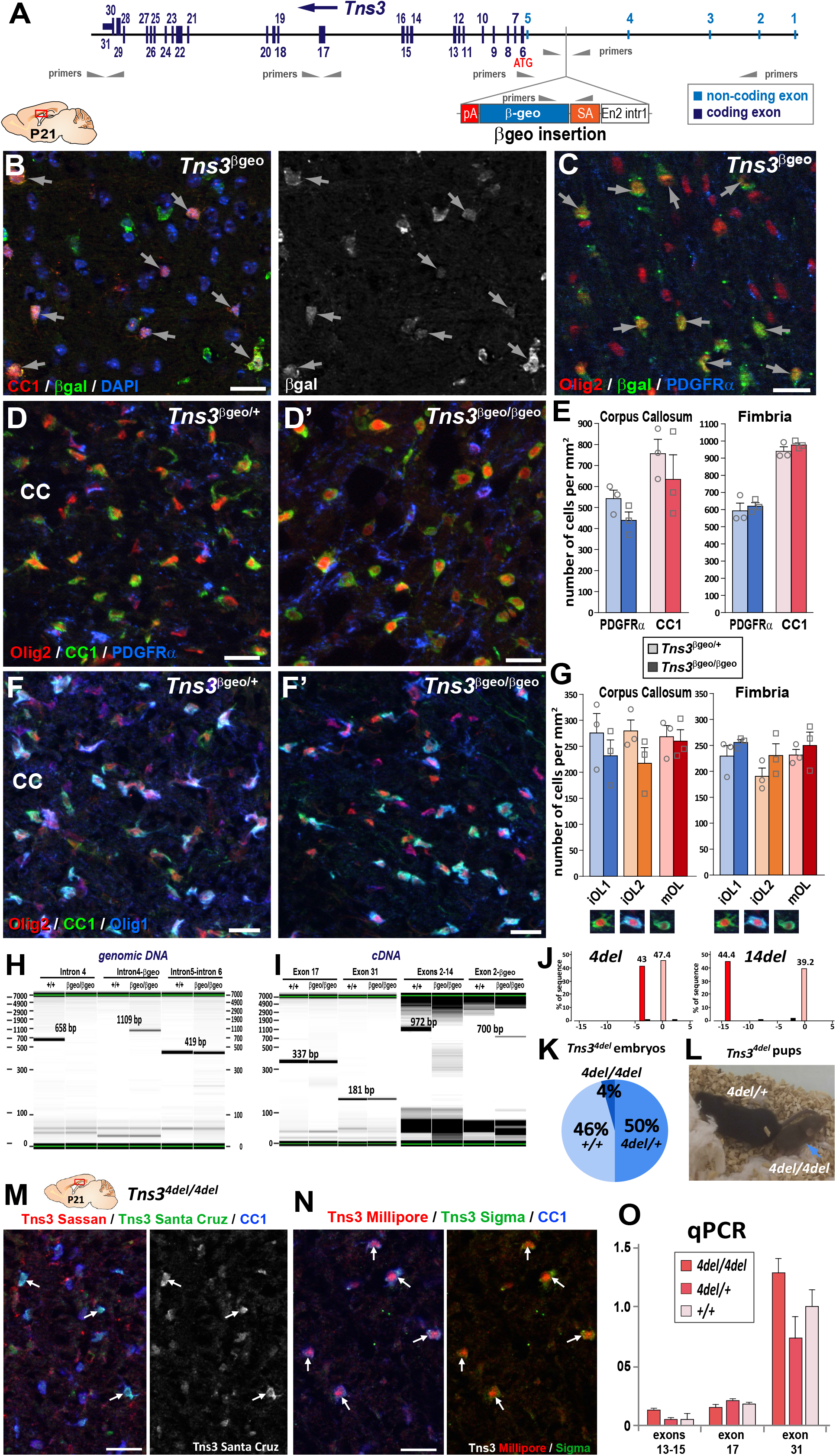
Oligodendrogenesis is normal in *Tns3* constitutive mutant mice, which still express *Tns3* full-length transcripts in the brain. **(A)** Schematic of *Tns3^βgeo^* allele, showing localization of primers used for PCR amplification. **(B,C)** β−galactosidase immunofluorescence in P21 sagittal brain sections, correspond to CC1^+^ OLs (arrows, B) or Olig2^high^/PDGFRα^−^ OLs (arrows, C) immature OLs. **(D,D’)** Immunofluorescence for PDGFRα^+^ OPCs and CC1^+^ OLs showing similar cell numbers in the CC of *Tns3^βgeo/+^*(D) and *Tns3^βgeo/βgeo^*(D’) brains. **(E)** Histograms representing density of PDGFRα^+^ OPCs and CC1^+^ OLs in the CC and the fimbria. (**F,F’**) Immunofluorescence for Olig2, CC1 and Olig1 allowing to identify three OL stages: CC1^high^/Olig1^-^ cells (named iOL1), CC1^high^/Olig1^high-cytoplasmic^ cells (iOL2) and CC1^low^/Olig1^cytoplasmic^ (mOLs), showing similar numbers of each OL stage in the CC of Tns*3^βgeo/+^* (F) and *Tns3^βgeo/βgeo^* (F’) brains. **(G)** Histograms representing the quantification of the density of iOL1s, iOL2s, and mOLs, in the CC and the fimbria. **(H)** PCR from genomic DNA showing the presence of the native Intron 4 only in wild type (+/+) mice, of the Intron 4-βgeo only in *Tns3^βgeo/βgeo^* mice, and of Intron 5 / Intron 6 in both mice. **(I)** Caliper visualization of PCR from cDNA showing the presence of Exon17 and Exon31 both in *Tns3^βgeo/βgeo^* and wild type mice, despite the amplification with primers for Exons 2-14 only in wild type mice, and with primers for Exon2-βgeo only in *Tns3^βgeo/βgeo^*mice. **(J)** Genotyping by TIDE analysis of *Tns3^4del^* and *Tns3^14del^* heterozygous mice showing wild type Tns3 allele and the 4 nucleotides deletion in *Tns3^4del^* allele and the 14 nucleotides deletion in *Tns3^14del^* allele. **(K)** Pie charts representing the genotypes of E14.5 embryos obtained from *Tns3^4del^* heterozygous inter-crosses. Note the sub-lethal phenotype indicated by the reduced number of *Tns3^4de/4del^* embryos compared to Mendelian ratios. **(L)** *Tns3^4del^* mice presenting growth defects in homozygous pups (arrow) at P14 compared to their heterozygous littermates. **(M,N)** Immunofluorescence showing that Tns3 is still detected in CC1+ OLs of *Tns3^4de/4del^* P21 mice, with four different Tns3 recognizing antibodies: (M) Sassan Hafizi lab and Santa Cruz, (N) Millipore and Sigma. **(O)** Histogram representing qPCR on cDNA from P21 brains showing no differences in the amplification of *Tns3* Exon13-15, Exon 17 and Exon31 between *Tns3^4del/4del^*, *Tns3^4del/+^* or WT mice, indicating the presence of a long *Tns3* transcript containing these exons in *Tns3^4del/4del^* brains. CC, corpus callosum. Scale bar, 20 μm.

**Figure S9.**
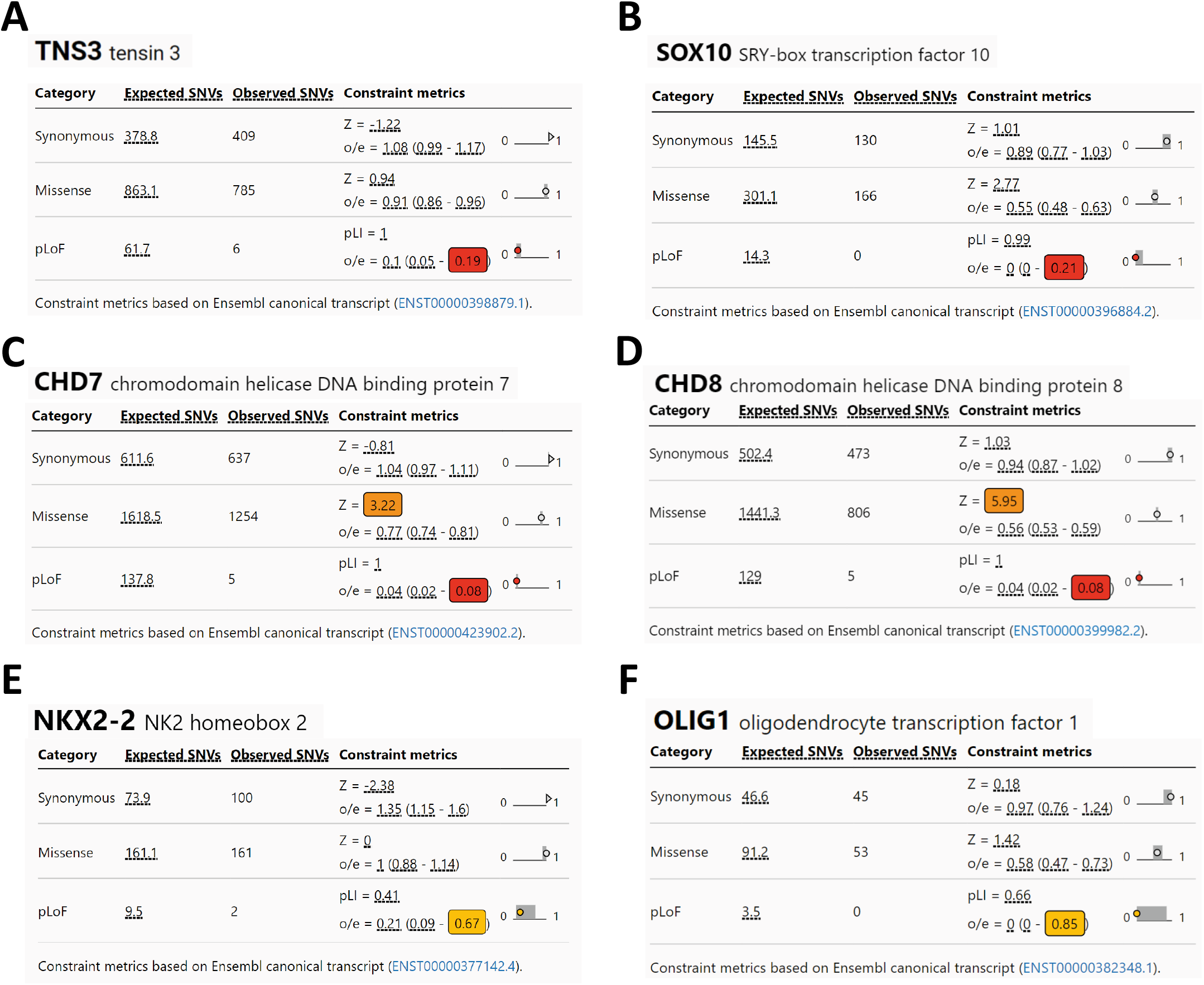
Intolerance for *TNS3* loss-of-function variants in the human population. **(A-F)** GnomAD data for *TNS3* and key regulators of oligodendrogenesis (*SOX10, CHD7, CHD8, NKX2-1*, and *OLIG1*) showing frequency and scores of different genetic variants in the human population: synonymous, missense, and pLOF (nonsense, splice acceptor, and splice donor variants). The numbers highlighted colored squares correspond to the LOEF (loss-of-function observed/expected upper bound fraction), that it is a conservative estimate of the observed/expected ratio. Low LOEUF scores indicate strong selection against predicted loss-of-function (pLoF) variation in a given gene, while high LOEUF scores suggest a relatively higher tolerance to inactivation. Note that *TNS3*, like *SOX10*, *CHD7*, and *CHD8* have very low LOEUF scores indicating high intolerance for their inactivation, contrary to *NKX2-1* and *OLIG1*, which are show more tolerance to their inactivation.

**Figure S10.**
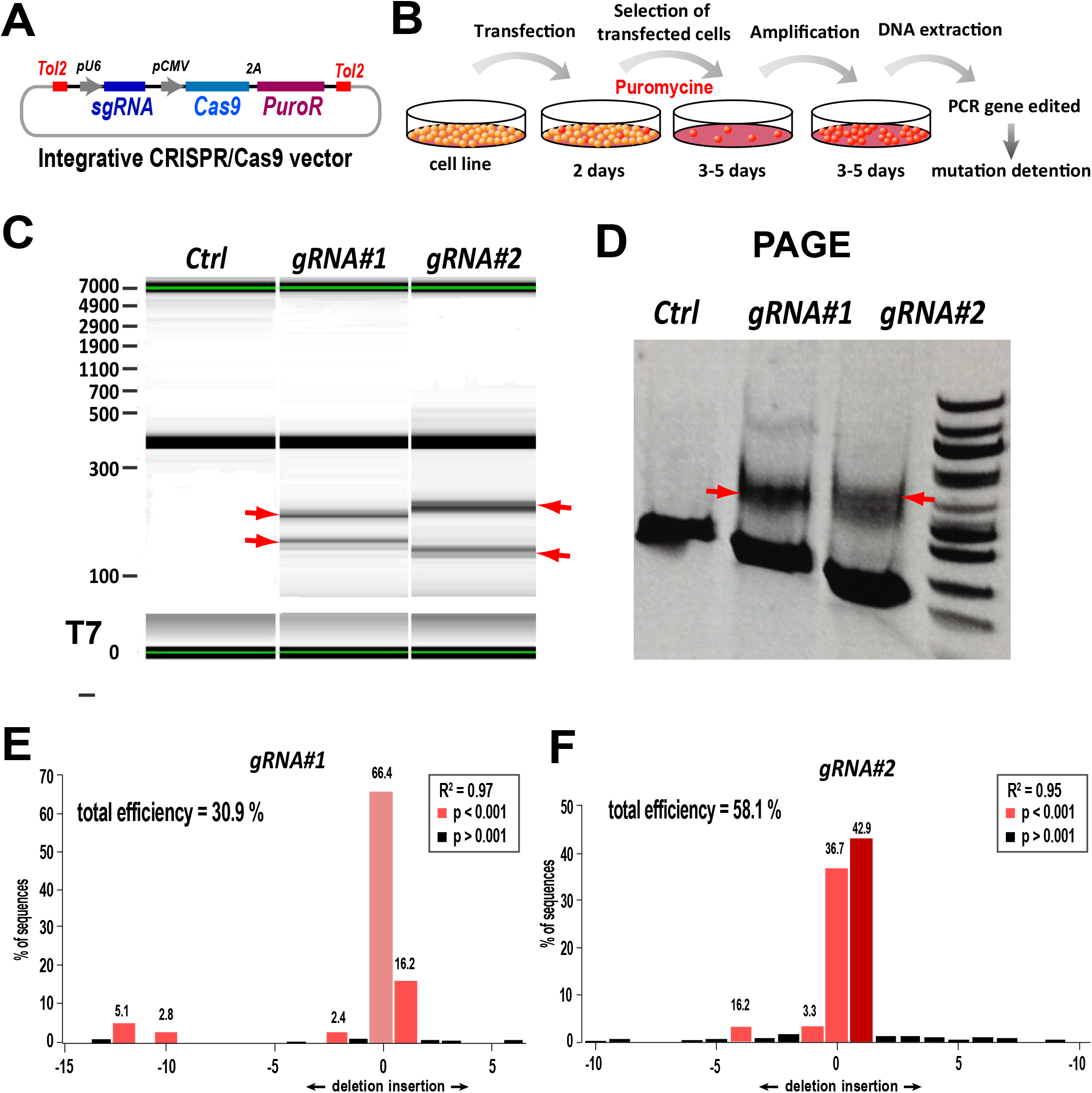
Generation and validation of *Tns3*-targeting CRISPR/Cas9 tools. **(A)** Schematic of the *Tol2-PX459* CRISPR/Cas9 expression vector allowing Tol2-DNA integration driving Cas9 & puromycin resistance expression (from polycistronic 2A-mediated cleaved) driven by CMV promoter and sgRNA expression from U6 promoter. **(B)** Scheme of C17.1 neuroblastoma cell line lipofection followed by puromycin selection of transfected cells used to amplify by PCR the *Tns3* targeted region and assess for indel mutations. **(C)** Caliper visualization of PCR products obtained after T7-mediated cleavage of DNA heteroduplex due to the indel mutations, showing smaller products (arrows) generated by Cas9 cutting with gRNA#1 or gRNA#2. **(D)**. Visualization Tns3-targetted region amplified by PCR run in a PAGE gel. Note the extra bands in gRNA#1 or gRNA#2 transfected cells corresponding to Cas9 cut products, contrary to *Tol2-PX459* empty plasmid (control). **(E,F)** TIDE analysis representing the percentage of sequences presenting a given indel mutation in cells transfected with gRNA#1 (E) or gRNA#2 (F).

**Figure S11.**
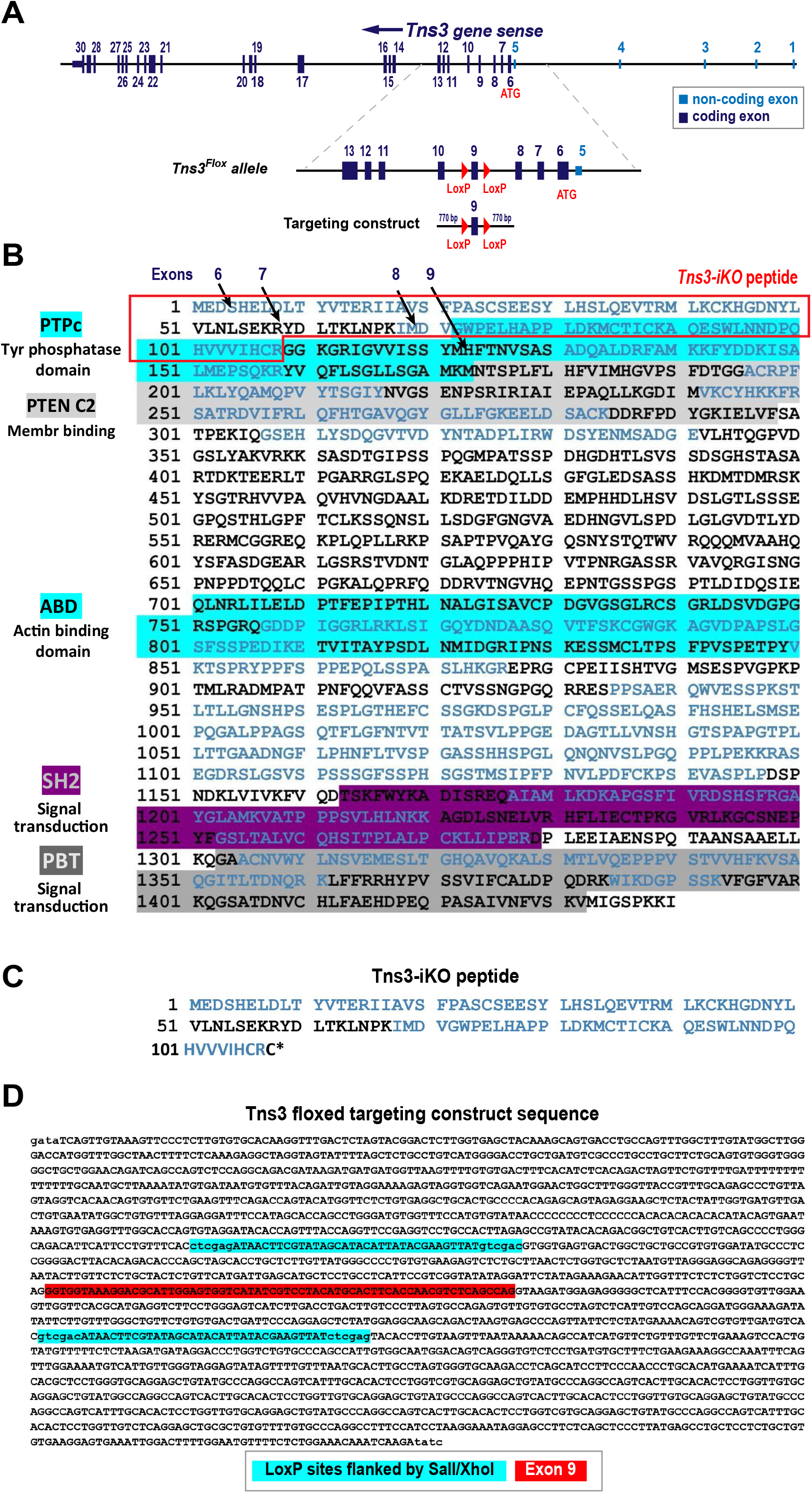
Generation of *Tns3* floxed allele. **(A)** Schematic of *Tns3* locus, *Tns3^Flox^*allele and targeting vector with LoxP sites flanking Exon9, used to induce homologous recombination in mouse ESCs. **(B)** Full-length Tns3 amino-acid sequence indicating its protein domains. Squared in red highlights the 109 amino-acid sequence of the Tns3 peptide produced upon Cre-mediated recombination in *Tns3^flox^* expressing cells. **(C)** Peptide sequence of protein product predicted for *Tns3-iKO* expressing cells. **(D)** *Tns3^Flox^* targeting vector sequence used for homologous recombination in mouse ESCs and generation of *Tns3^Flox^*mice, with LoxP sites highlighted in blue and Exon 9 highlighted in red.

**Figure S12.**
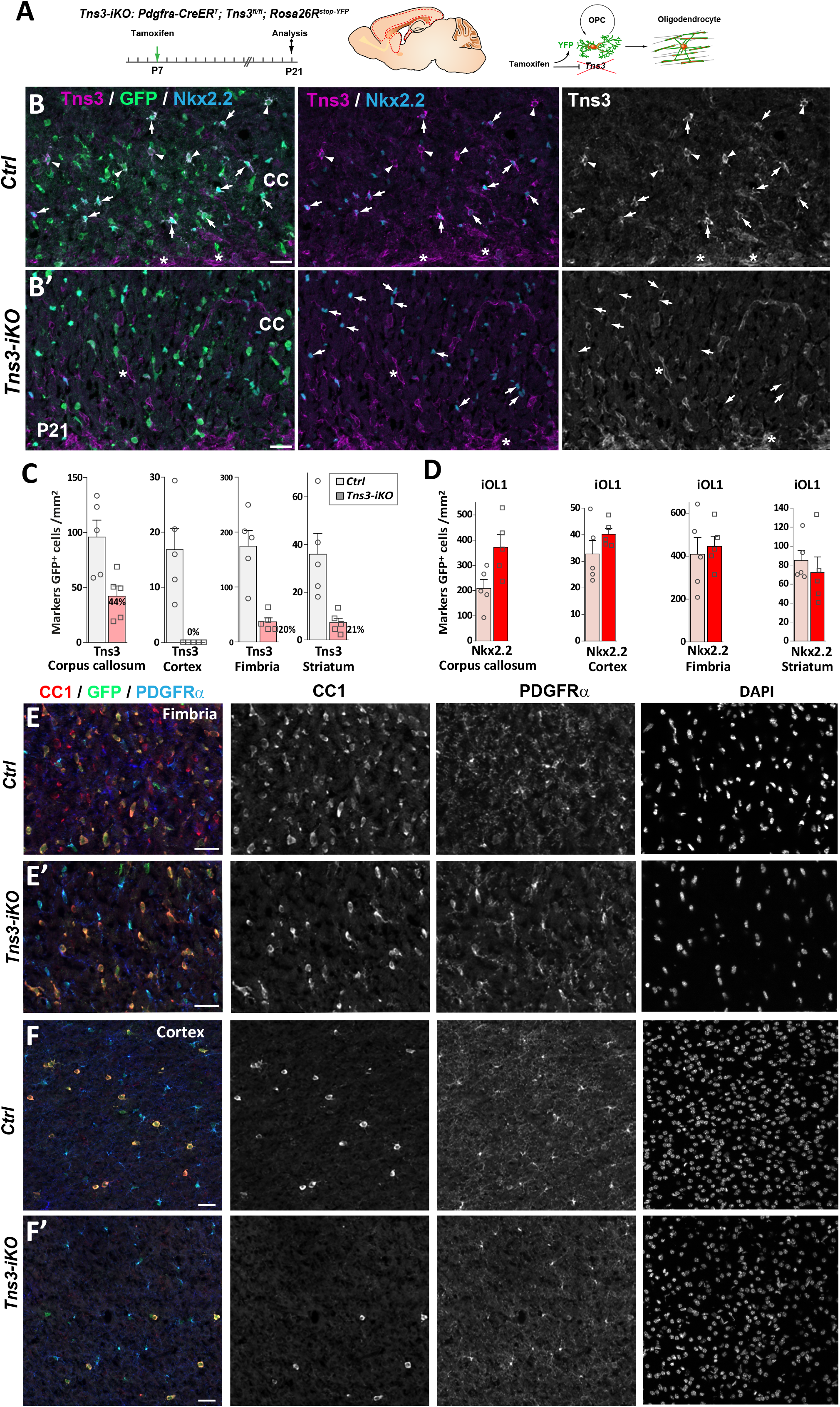
Efficient OPC-specific *Tns3* deletion in the postnatal brain and reduced oligodendrocyte generation from OPCs. **(A)** Scheme of tamoxifen administration to *Tns3-iKO* (*Cre^+^; Tns3^flfl^*) and control (*Cre^+^; Tns3^+/+^*) mice, Cre-mediated genetic changes, and timing of experimental analysis. **(B-B’)** Immunofluorescence for Tns3, GFP and Nkx2.2 in P21 mice sagittal brain sections at the level of the corpus callosum, illustrating the loss of Tns3 signal in Nkx2.2^high^ iOL1s (arrows) but not in vessels (asterisk) of *Tns3-iKO* (B’) compared to control (B). Note that Nkx2.2^high^ iOL1s do not change in density in *Tns3-iKO* compared to control. **(C)** Histograms showing the Tns3+ cells density in the corpus callosum, fimbria, cortex, and striatum, in *Tns3-iKO* and control. **(D)** Histograms showing the Nkx2.2^high^ iOL1s density in the corpus callosum, fimbria, cortex, and striatum, in *Tns3-iKO* and control. **(E-F’)** Immunostaining corresponding to Fig.4B-C’ panels showing individual channels for CC1, PDGFRα, and DAPI in the fimbria (E,E’) and cortex (F,F’) of control mice (E,F) and *Tns3-iKO* mice (E’,F’). CC, corpus callosum. Scale bar, 20 μm.

**Figure S13.**
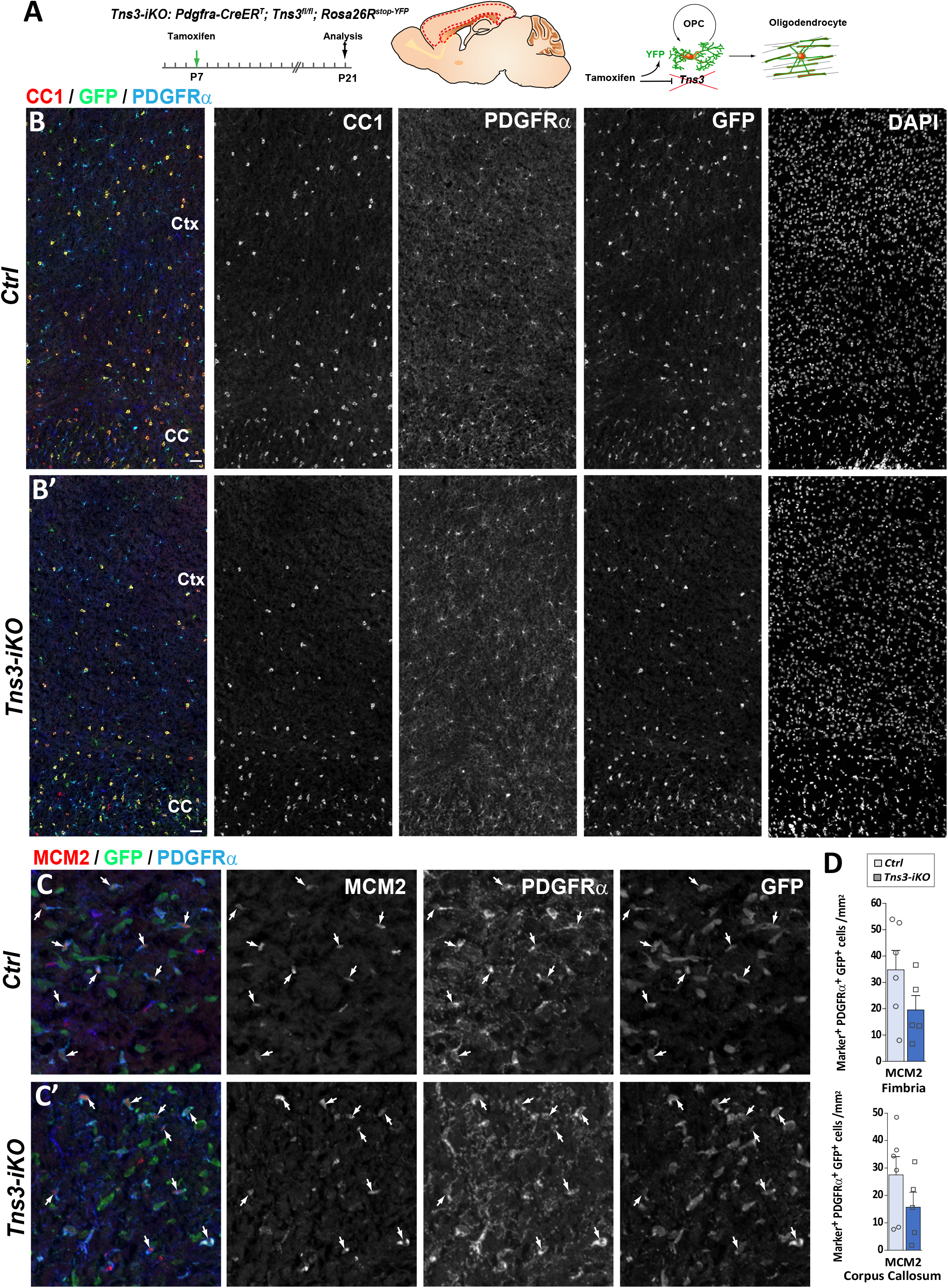
OPC-specific *Tns3* deletion reduces oligodendrocyte generation without changing OPC proliferation in the postnatal brain. **(A)** Scheme of tamoxifen administration to *Tns3-iKO* (*Cre^+^; Tns3^flfl^*) and control (*Cre^+^; Tns3^+/+^*) mice, Cre-mediated genetic changes, and timing of experimental analysis. **(B-B’)** Immunostaining illustrating at low magnification the reduction in number of CC1^+^ OLs but not PDGFRα^+^ OPCs in the cortex and corpus callosum of *Tns3-iKO* mice (B’) compared to control mice (B), showing each individual channel and DAPI. **(C-C’)**. Immunofluorescence for MCM2 proliferative marker, GFP, and PDGFRα illustrating similar proportion of proliferative OPCs in control (C) and *Tns3-iKO* mice (C’). **(D)** Histograms quantifying the proliferative fraction of GFP^+^ OPCs in the fimbria (Upper) and corpus callosum (Lower). Note the slight reduction (non-significant) in proliferation in *Tns3-iKO* OPCs compared to control OPCs. CC, corpus callosum; Ctx, cortex. Scale bar, 20 μm.

**Figure S14.**
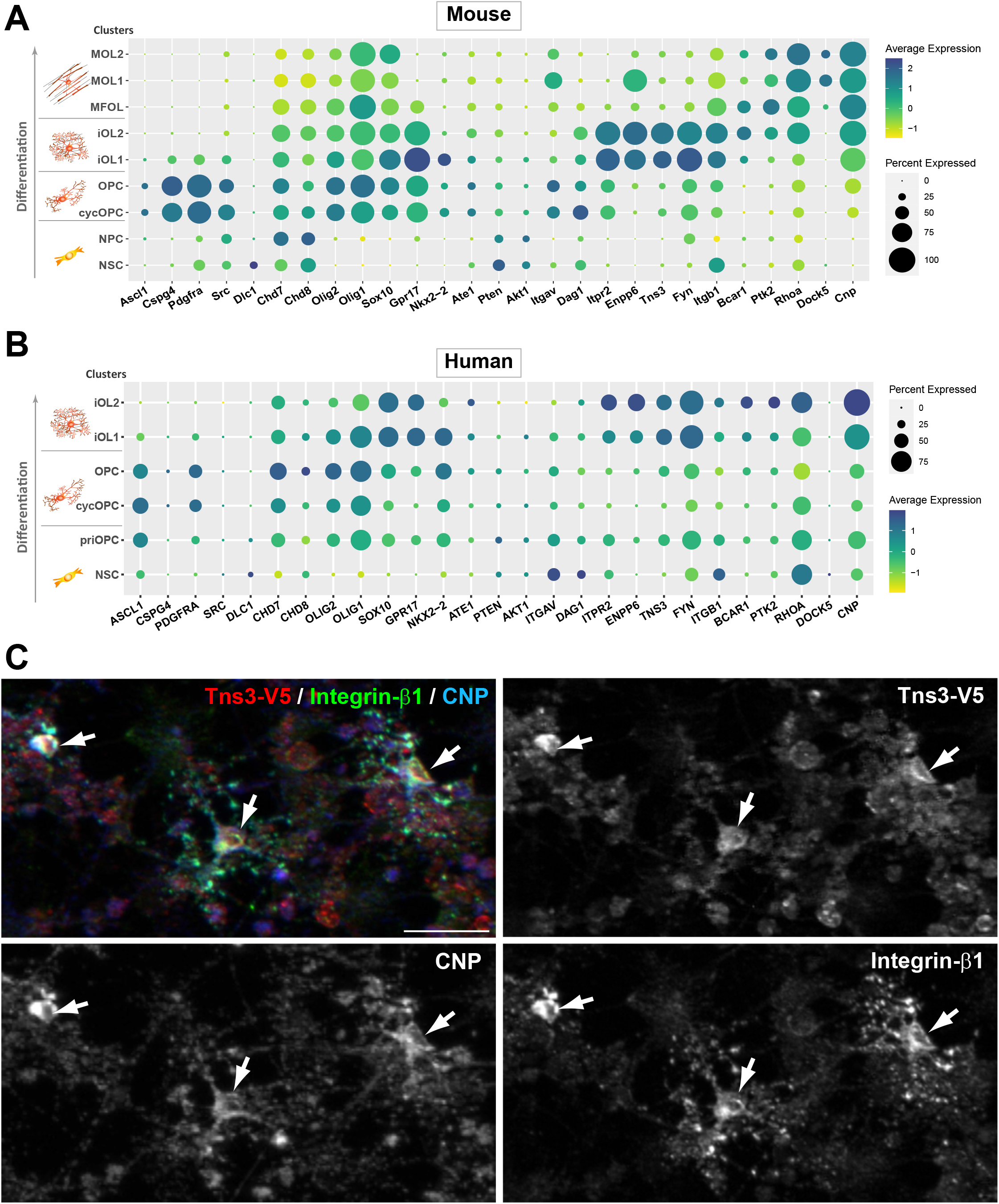
*Tns3* is co-expressed with integrin signaling genes in immature oligodendrocytes. **(A,B)** Dotplots showing *Tns3/TNS3* expression patter in mouse (A) and human (B) single cells at different stages of oligodendrogenesis, together with key markers of each cluster and genes involved in integrin signaling pathway. Note that high levels of Tns3/TNS3 in immature oligodendroctyes (iOL1 and iOL2) together with known genes of integrin pathway previously implicated in oligodendrogenesis (*Itgb1, Fyn, FAK/Bcar1, p130Cas/Ptk2*) both in mouse and human cells. **(C)** Immunofluorescence in neurosphere-derived neural progenitor cultures generated from *Tns3^Tns3-V5^* neonatal brain after 3 days of differentiation showing co-expression of Tns3-V5 (V5 antibody) and Integrin-β1 in CNP^+^ differentiating OLs (arrows). Scale bar, 20 μm.

**Figure S15.**
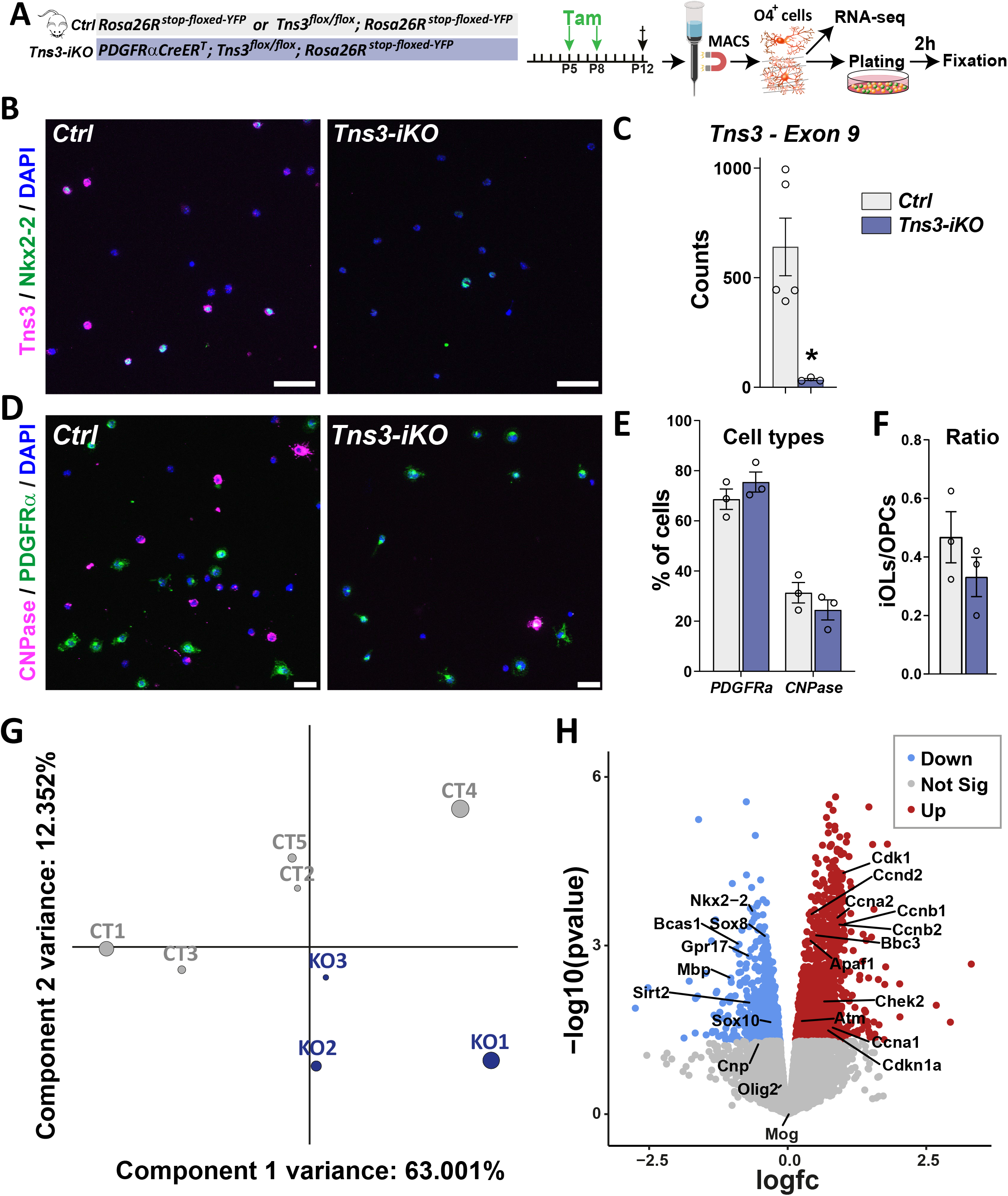
Transcriptomic analysis of acutely Tns3-deleted oligodendroglia. **(A)** Diagram representing tamoxifen (Tam) injection at P5 and P8 Ctrl and *Tns3-iKO* pups followed by the MACSorting of O4^+^ cells used for RNAseq and plating. Cells were fixed 2h after and used to do immunstaining. **(B)** Immunostaining validating the loss of Tns3 protein in *Tns3-iKO* O4^+^ cells compared to Ctrl. **(C)** Graph representing the count of exon 9 of Tns3 in the RNAseq data. Note the reduction of floxed exon in the *Tns3-iKO* O4^+^ cells compared to control (*Ctrl*). **(D)** Immunostaining showing the composition in OPCs (PDGFRα^+^ cells) and iOLs (CNPase^+^ cells) among *Ctrl* and *Tns3-iKO* O4^+^ cells after MACS. **(E)** Graph indicating that the same amount of OPCs (PDGFRα^+^ cells) and iOLs (CNPase^+^ cells) were found in *Ctrl* and *Tns3-iKO* O4+ cells after MACS. **(F)** Graph showing that the ratio of iOLs (CNPase^+^ cells)/OPCs(PDGFRα^+^ cells) was not altered in *Tns3-iKO* O4^+^ cells compared to Ctrl. **(G)** Principal component analysis (PCA) of the RNA-seq data transcripts of *Ctrl* (CT) and *Tns3-iKO* (KO) O4^+^ cells. **(H)** Volcano plot showing the upregulated (red) and downregulated (blue) in *Tns3-iKO* O4^+^ cells compared to *Ctrl*. Some example of genes can be found. Scale bars: B, 50 μm; D, 10 μm.

## Peer-reviewing process

We thank all reviewers for their positive and encouraging comments and criticisms to improve our work. Here we present a reviewed version of the manuscript according to the comments risen.

*Reviewer #1 (Evidence, reproducibility and clarity (Required)):*

*This is an interesting paper that identifies Tns3 as a potential effector of oligodendrocytes differentiation based on an ingenious strategy comparing regulatory binding sites of known master regulators of differentiation, and then shows using in vivo genetics that this role is indeed correct. Next, a potential mechanism is identified by showing co-localization with beta 1 integrin, known to regulate apoptosis of newly-formed oligodendrocytes. The results are well illustrated and the experiments performed with appropriate power using a broad range of techniques that combine in silico, in vitro and in vivo work to great effect*.

*I think this represents an important contribution that will be of significant interest to neuroscientists - the mechanisms regulating oligodendrocytes generation remain poorly understood and the evidence that this contributes to adult learning (adaptive myelination) and CNS regeneration makes this a key question. I would suggest that the following are considered before publication:*

We thank the reviewer for this positive comments and critics to improve the manuscript.

*The work describing the KO mice that were not used as they proved unsuitable need not be described - it breaks the logical flow*.

In agreement with the reviewer comment, we have reduced this part to a sort paragraph indicating that our analyses of several Tns3 constitutive KO lines showed developmental lethality and possible genetic compensation in Tns3 expression, leading us to conclude them inappropriate tools to study Tns3 function in oligodendrogenesis. We have summarized the data in Fig. S7 and the description in the method section.

*It would be useful to compare the extent of cell death in the Tns3 cKO mice with that described in the alpha6 integrin KO and the integrin beta1 cKO (the Colognato and Benninger papers). Do they match? If not (and I suspect the Tns3 cKO death is greater) could other mechanisms be downstream of the Tns3?*

In agreement with the reviewer comment, we have added the following paragraph to the discussion:

‘Knockout mice for *integrin-α6* present a 50% reduction in brainstem MBP^+^ OLs at E18.5, just before they die at birth, accompanied by an increase in TUNEL^+^ dying OLs (Colognato et al, 2002). Similarly, conditional deletion of *integrin-β1* in immature OLs by *Cnp-Cre* also leads to a 50% reduction in cerebellar OLs at P5, with a parallel increase in TUNEL^+^ dying OLs (Benninger et al., 2006). Therefore, given that *Tns3*-induced deletion in postnatal OPCs also leads to 40-50% reduction in OLs in both grey and white matter regions of the postnatal telencephalon (this study), paralleled by similar increase in TUNEL^+^ apoptotic oligodendroglia, we suggest that Tns3 is required for integrin-β1 mediated survival signal in immature oligodendrocytes.’

*I’m not sure why the authors argue that the activation of beta 1 would not be informative experiment? This will regulate actin dynamics just as it regulates other integrin signaling pathways. Indeed, I would argue that an integrin activation experiments would be a neat way to prove mechanism (as it would be predicted to rescue the Tns3 cKO phenotype)*.

In agreement with the reviewer comment, we have removed this sentence: ‘If so, exogenous activation of integrin α6β1 in cultured OPCs by Mn2+ (Colognato et al., 2004) would not be expected to increase oligodendrogenesis in Tns3-iKO oligodendroglia.’

In an effort, to understand Tns3 function by acute Tns3-deletion in postnatal OPCs, we have compared the transcriptome of *Tns3-iKO* oligodendroglia compared to control cells, and we present these results in figure 7 pinpointing deregulated genes leading to reduced oligodendroglial differentiation, **integrin dysregulation**, increase apoptosis, and conflicting cell cycle signaling, and leaving for further studies the full characterization how the loss of Tns3 leads to the deregulation of these processes.

*Can the authors provide any data on GM oligos and their OPCs? Is the requirement for Tns3 the same, and if so what might the implications be in the adult where new oligodendrocytes are being generated throughout life?*

Indeed, in our analyses of *Tns3-iKO* mice, we provide quantifications of the cortex as a grey matter territory, showing a similar 40-50% reduction in OLs as in white matter areas (corpus callosum and fimbria, and mixed regions such as the striatum.

*I note in S13 that integrin beta1 is not highly expressed in human oligos at the time in question. Does this call into question the relevance for human disease?*

We realize that scRNAseq plots are never easy to interpret but it is important to note that the levels of expression are coded by the intensity of the color scale, while the surface of the dot plots indicate the experimental sensitivity to detect transcript expression in a larger or smaller proportion of the cells in a given cluster/cell type (due to the drop out limitation of current single cell RNA-seq technologies). Considering this, please note that beyond a stronger expression in neural progenitor cells (NPCs, blue color), integrin-b1 (*Itgb1*) transcripts are expressed at medium to high levels (green to blue) in human immature OLs (Fig. S13B), similar to their pattern of expression in mouse oligodendroglia (Fig. S13A).

Reviewer #1 (Significance (Required)):

See above

Reviewer #2 (Evidence, reproducibility and clarity (Required)):

*In this article, the authors identify and characterise Tensin3 (Tns3) as a target of key oligodendroglial transcription factors driving differentiation in the mouse. They use multiple transgenic models to describe loss of function, and suggest Tns3’s action through integrin B1 signalling, with the key function being oligodendroglial survival*.

*There is extensive and impressive work here, including identification of Tns3 by ChIPseq, expression of Tns3 in brain development, analysis of human (ES-derived) and mouse scRNAseq to infer timing of expression in the differentiation pathway, generation of V5-tagged Tns3-KI mice to overcome antibody limitations, identification of its expression in mouse remyelination, generation of a new Tns3KO mouse, in vivo Crispr Tns3KO in development, generation of a conditional KO, for deletion in adulthood, and finally some culture work to investigate potential mechanisms of actions. The bottom line is that Tns3 is required for survival of OPCs and immature oligodendrocytes in development/remyelination in mouse at least, and loss leads to apoptosis (through p53 increase and loss of integrin-B1 signalling), leading to a failure of proper differentiation*.

*The experiments are carefully done, convincing and the tools generated impressive. There is clearly more to be done on clarifying the mechanism of action of Tns3, but I do not think further experiments on this topic are needed for this paper - they can wait for the next*.

We thank the reviewer for the positive and encouraging reviewing comments. In an effort, to understand Tns3 function by acute Tns3-deletion in postnatal OPCs, we have compared the transcriptome of *Tns3-iKO* oligodendroglia compared to control cells, and we present these results in figure 7 pinpointing deregulated genes leading to reduced oligodendroglial differentiation, integrin dysregulation, increase apoptosis, and conflicting cell cycle signaling, and leaving for further studies the full characterization how the loss of Tns3 leads to the deregulation of these processes.

*My only query is whether the expression of Tns3 is also in immature OLs in human brain (rather than human ES-derived OLs). This should be easily checked with interrogation of online Shiny apps from already published snRNAseq from various groups on human post mortem adult brain, but if not present then in also baby/fetal brain. This would be interesting and may well be different from the ES_derived cells which tend to be very immature and would add interest to the possible translational impact*.

According to the suggestion of the reviewer, we analyzed 69,174 snRNAseq GW9-GW22 from fetal cerebellum,; w, 2021; https://doi-org.proxy.insermbiblio.inist.fr/10.1038/s41593-021-00872-y), which we present now in Figure S3, finding a cluster of cells expressing iOL markers, including *NKX2-2, TNS3, ITPR2*, and *BCAS1*, similar to the hiPSCs-derived iOL1/iOL2 clusters and mouse iOL1/iOL2 clusters shown in Fig. S2.

We also analyzed other datasets without finding iOLs given their age or numbers, including:

- Immunopanned PDGFRA+ cells from human cortex GW20-GW24 (2690 cells, Huang and Kriegstein, Cell 2020) finding OPCs but not iOLs.

-The recently published dataset from GW8-GW10 human forebrain oligodendroglia (van Brugen & Castelo-Branco, Dev Cell 2022; https://doi.org/10.1016/j.devcel.2022.04.016) containing OPCs but not iOLs.

-The GW17 to GW18 human cortex (40,000 cells, Polioudakis & Geschwind, 2019, https://doi.org/10.1016/j.neuron.2019.06.011) containing OPCs but not iOLs.

*Reviewer #2 (Significance (Required)):*

*This work extends our knowledge of oligodendroglial differentiation, links it to the ECM and provides interest in manipulating this in diseases including glioma*.

*My expertise: myelin, oligodendroglia, remyelination, human neuropathology*

*Reviewer #3 (Evidence, reproducibility and clarity (Required)):*

*see below*

*Reviewer #3 (Significance (Required)):*

*Using purified oligodendrocytes target genes of key regulators of oligodendrocyte differentiation were analyzed, which led to the identification of Tensin-3. The authors performed a detail characterization of Tensin-3 expression. They found that Tensin-3 is highly expressed in immature mouse and human oligodendrocytes. Interestingly, Tensin-3 is selectively enriched in immature oligodendrocytes, and not present at detectable levels in OPCs and mature oligodendrocytes. Subsequently, the authors characterized Tensin-3 function by a series of knockdown approaches in vitro and in vivo. These series of experiments revealed an essential function of Tensin-3 in supporting oligodendrocytes survival. In the absence of Tensin-3 a large fraction of oligodendrocytes undergo apoptosis while differentiating to mature oligodendrocytes*.

*This is a remarkable study applying an impressive array of methods that led to an important discovery in the field of oligodendrocyte biology. The main advances for the field are: 1) identification of a novel marker for premyelinating oligodendrocytes, 2) elucidation of Tensin-3 as a pro-survival factor in oligodendrocytes differentiation, 3) evidence of link of Tensin-3-integrin signal in survival of oligodendrocytes*.

*The data is well presented and organized, and the paper well written*.

*I recommend publication with only minor suggestions for a revision:*

We thank the reviewer for this positive comments and critics to improve the manuscript.

*In Figure 2, only images are shown, and the data is referred to as highly expressed or strong co-localization. Even if the data looks clear, the authors should provide some quantification of the data in the figure*.

We thank the reviewer for his comment and we have now provided a quantification of the fraction of Tns3+ cells expressing different markers of oligodendrocyte lineage progression/stages, and the percentage of each stage expressing Tns3.

Figure 3 is given too much weight in the manuscript text. I would recommend to shorten the text in the result section, and to move this figure to the supplement as it does not advance the story. It mainly shows that the KO mice still express transcripts in the brain. Were the transcripts lost in peripheral tissue?

As mentioned above, in agreement with the reviewers #1 and #3 comments, we have reduced this part to a sort paragraph indicating that our analyses of several Tns3 constitutive KO lines showed developmental lethality and possible genetic compensation in Tns3 expression, leading us to conclude them inappropriate tools to study Tns3 function in oligodendrogenesis. We have summarized the data in Fig. S7 and the description in the method section.

*Page 11: the authors describe in the text how the floxed allele was generated. This should be shifted to the supplement*.

According to reviewers suggestion, we have moved the description of Tns3 floxed allele generation to the Methods section.

*Page 16: the authors refer to Bcas1 as a problematic marker for immature oligodendrocytes, because the transcript is also expressed in mature oligodendrocytes. The authors are correct that the transcript is expressed in mature oligodendrocytes. However, the proteins changes its localization when oligodendrocytes mature. On protein level, it is valuable and a selective marker, as antibodies only label pre-myelinating and actively myelinating cells. In mature oligodendrocytes, antibodies against Bcas1 do not label the cell, only myelin. The text is misleading and needs to be corrected*.

In agreement with reviewers comment we have modified the text as follows: ‘An optimized protocol for immunodetection using Bcas1-recognizing antibodies has been shown to label iOLs (Fard et al., 2017).’

